# Primitive steroidogenesis in mast cells: A novel regulatory mechanism for mast cell function

**DOI:** 10.1101/2025.02.05.636621

**Authors:** Jhuma Pramanik, Qiuchen Zhao, Yumi Yamashita-Kanemaru, Hosni Hussein, Natalie Z. M. Homer, Soura Chakraborty, Sanu K. Shaji, Klaus Okkenhaug, Rahul Roychoudhuri, Bidesh Mahata

## Abstract

Mast cells, ancient immune sentinels, are crucial in immune responses, tissue homeostasis and inflammatory pathologies. This study unveils a previously unknown regulatory mechanism in mast cell biology: *de novo* steroidogenesis. Through comprehensive multi-omics analysis and functional assays, we demonstrate that mast cells express Cyp11a1 and produce pregnenolone, representing a primitive form of steroidogenesis. This cell-intrinsic steroidogenic pathway is essential for mast cell development, survival, and functional regulation. Genetic abrogation of mast cell steroidogenesis leads to exaggerated inflammatory and anaphylactic responses *in vivo*. Our integrative approach reveals extensive transcriptional and proteomic remodelling during mast cell regranulation, with steroidogenesis playing a pivotal role in coordinating recovery and tissue repair processes. We uncover significant sexual dimorphism in mast cell proteomes and a global uncoupling of transcriptional and translational programmes. These findings advance our understanding of mast cell physiology and provide a foundation for developing targeted therapies for mast cell-associated pathologies.

## INTRODUCTION

Mast cells are ancient immune sentinels of hematopoietic origin strategically positioned at external and internal barrier tissues (Dahlin et al., 2022; Galli et al., 2020; Galli et al., 2005; Gentek et al., 2018; Gurish and Austen, 2012; Voehringer, 2013). These evolutionarily conserved cells are found in all vertebrates and predate the emergence of adaptive immunity, with their origins traced back to urochordates (Crivellato and Ribatti, 2010; St John et al., 2023). Mast cells are crucial in initiating and orchestrating immune responses, maintaining vascular homeostasis, and contributing to tissue remodelling (Galli et al., 2020; Galli et al., 2005; Voehringer, 2013). Two major mast cell types have been identified in rodents and humans: connective tissue mast cells and mucosal mast cells (Dahlin et al., 2022; Galli et al., 2020; Galli et al., 2005; Gentek et al., 2018; Gurish and Austen, 2012; Voehringer, 2013). These cells are characterized by their dense secretory granules containing preformed mediators such as histamine, tissue necrotic factors, serotonin, and a diverse array of mast cell-specific proteases (Wernersson and Pejler, 2014). Upon activation, mast cells rapidly release these granule contents, followed by the synthesis and secretion of pro-inflammatory lipid mediators, chemokines, and cytokines (Wernersson and Pejler, 2014). However, these long-lived cells can regranulate to re-acquire the secretory potential (Iskarpatyoti et al., 2022). What happens during the regranulation phase, reacquiring the secretory potential and for the restoration of immune and tissue homeostasis, is immensely interesting in the field.

Despite their critical physiological functions, dysregulation of mast cells can lead to various pathological conditions, including allergic and inflammatory disorders. While significant progress has been made in understanding mast cell biology, many aspects of their regulation remain elusive. Most known regulatory mechanisms fine-tuning mast cell function are associated with IgE-dependent pathways, primarily through inhibitory receptors containing immunoreceptor tyrosine-based inhibitory motifs (ITIMs) (Bulfone-Paus et al., 2017; Galli et al., 2008). However, given that mast cells evolved hundreds of millions of years before the emergence of adaptive immunity and IgE-dependent responses, it is likely that more primitive, IgE-independent regulatory mechanisms exist (St John et al., 2023).

Steroidogenesis is a biosynthetic process by which steroid hormones are made from cholesterol. CYP11A1 (also known as P450scc) catalyses the first and rate-limiting step of the steroidogenesis pathway and converts cholesterol to pregnenolone (Figure 1A, left panel) (Miller, 2017; Miller and Auchus, 2011). This biochemical step is also the most primitive step of the pathway. Pregnenolone is, therefore, the first bioactive steroid hormone of the pathway and precursor of all other steroid hormones. Steroid glands, such as the adrenal gland, ovary, testis and placenta, produce gland-specific steroid hormones to control several physiological processes in vertebrates, including salt balance, carbohydrate metabolism, sex differentiation and immune regulation (Miller, 2017; Miller and Auchus, 2011). While the regulation of mast cell function by steroid hormones is well-established and has been leveraged to treat various mast cell-associated pathologies (Oppong et al., 2013; Quatrini et al., 2021; Strehl et al., 2019), the reverse relationship has received no attention. The capacity of mast cells themselves to synthesize and secrete steroids remains unexplored. To date, no research has directly investigated mast cell-mediated steroidogenesis or its potential role in regulating biological processes, representing a significant gap in our understanding of mast cell biology and immune regulation.

**Figure 1.**
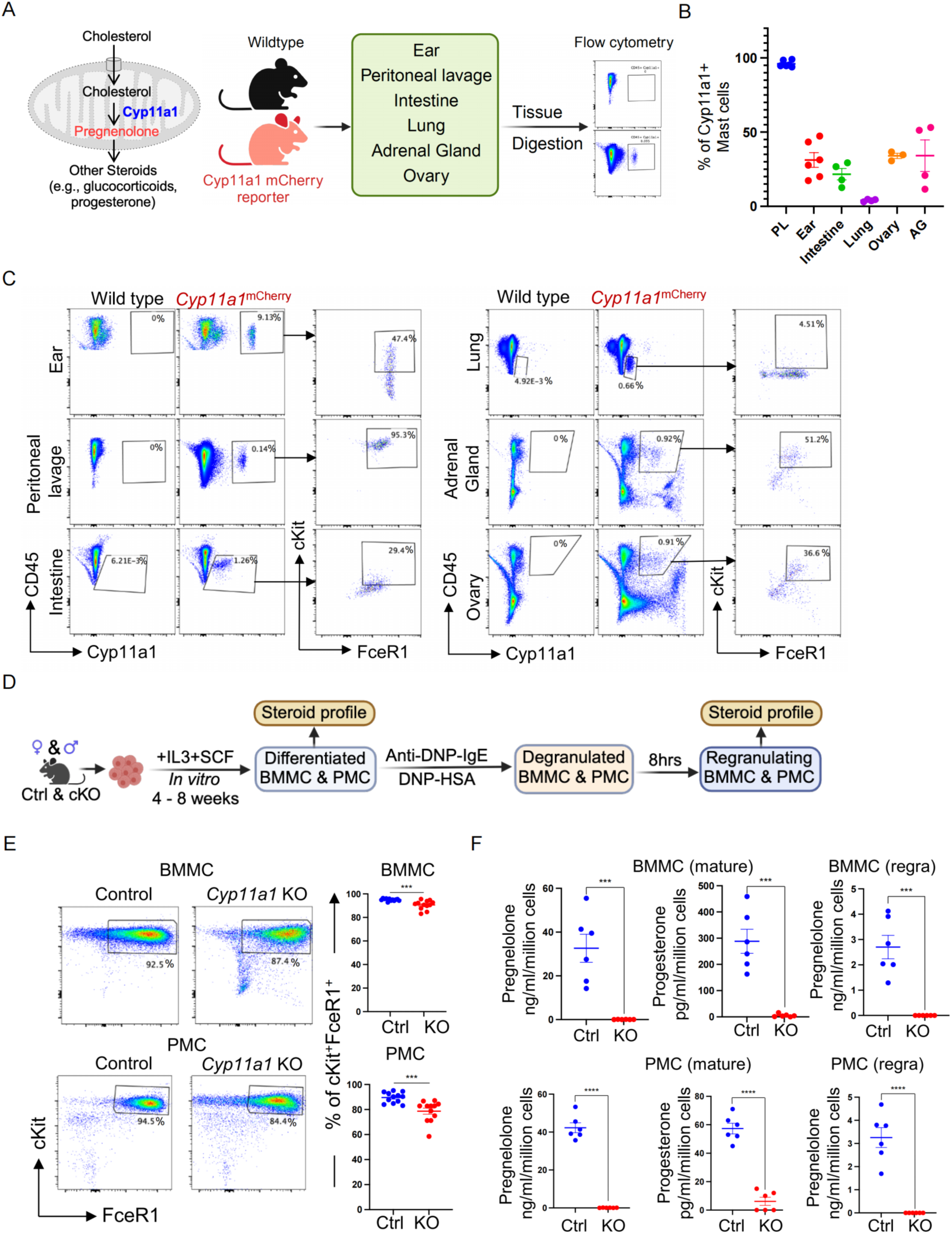
Cyp11a1 expression and steroid production by mast cells. **A.** A schematic representation of the Cyp11a1 activity catalysing the rate-limiting step of steroidogenesis (schematic on the left) and experimental plan (schematic on the right) used in B and C. Tissues and organs of healthy, unchallenged Cyp11a1-mCherry reporter or wild-type mice were mechanically dissociated and enzymatically digested. Single-cell suspensions were analysed by flow cytometry to measure Cyp11a1 expression. **B.** Graphical presentation of the Cyp11a1-mCherry expression in mast cells obtained from specified tissue origin. PL: Peritoneal lavage; AG: Adrenal gland. Representative of three independent experiments; each experiment contains 3–6 mice. **C.** Flow cytometric display of Cyp11a1-mCherry expression in mast cells (representative FACS plots). Representative FACS profile of Cyp11a1-mCherry expression shown in B. FACS gating: All cells > Singlets > live CD45^+^ cells > Lineage^-^ > Cyp11a1 mCherry^+^ cells > cKit^+^FceR1^+^ **D.** Schematic representation of the experimental plan used in E and F. *Cyp11a1*^fl/fl^ (Ctrl) or *Cyp11a1* ^fl/fl^ *Vav1*^Cre^ (KO) mice’s bone marrow or peritoneal cavity-derived mast cells (BMMC and PMC) were *in vitro* differentiated to obtain mature mast cells. Mature mast cells were sensitised by anti-DNP IgE antibody and activated by DNP-HSA. Eight hours after degranulation, cells were considered as regranulating mast cells. Cell culture supernatants of mature and regranulating mast cells were analysed by LC-MS/MS for the presence of all steroid hormones (steroid profiling); and mature mast cells were analysed by flow cytometry. **E.** Effect of *Cyp11a1* knockout in bone marrow-derived (BMMC) and peritoneal cavity-derived (PMC) mast cell maturation (as defined by cKit and FceR1 expression). Left panel: representative FACS plot. Right panel: Graphical presentation of cKit^+^FceR1^+^ cells. Representative of five independent experiments. **F.** Steroid profiling of mature and regranulating mast cell (BMMC and PMC) culture supernatant. Only those steroids that show Cyp11a1-dependent synthesis, pregnenolone and progesterone, are displayed. Mature mast cells were cultured for 4 days, and regranulating mast cells were cultured for 8 hours. (N=6). p-value was calculated using unpaired two-tailed t-test.

Recent advancements in computational biology and molecular detection methods, particularly omics approaches, have revolutionized our understanding of cellular processes (Dai and Shen, 2022). Comprehensive transcriptomics analyses, at single-cell or population level, have revealed the heterogeneity of mast cell populations and their developmental origins (Dahlin et al., 2022; Derakhshan et al., 2022; Dwyer et al., 2016; Tauber et al., 2023). Proteomic studies have provided insights into the mast cell protein repertoire of released proteins, phosphoproteins, and surface glycoproteins (Cao et al., 2007; Freitas Filho et al., 2019; Gschwandtner et al., 2017; Plum et al., 2020; Saha et al., 2022; Shubin et al., 2017). However, an integrative large-scale omics approach encompassing transcriptomics, proteomics, and steroidomics to elucidate mast cell biology across different functional states, tissue origins, and genders remains to be conducted.

In this study, we employ an integrative omics approach of transcriptomics, proteomics and steroidomics to visualise the in-depth repertoire of regulatory factors in cell-intrinsic regulation of mast cell maturation (i.e., differentiation from progenitors or precursors), activation and regranulation (after degranulation), their tissue origin and gender-specific differences. We discovered that mast cells express Cyp11a1 and produce steroid pregnenolone, representing a primitive form of steroidogenesis. Cyp11a1 expression and pregnenolone synthesis are elevated during the re-granulation phase. Mast cell steroidogenesis is required to regulate their inflammatory function negatively, possibly for fine-tuning their effector function. Under homeostasis, however, it is required for their survival and maturation. In the absence of mast cell steroidogenesis, mice exhibit overt allergic and anaphylactic responses in the experimental anaphylaxis model. Through comprehensive transcriptome and proteome analyses, we uncover the unprecedented cell-intrinsic role of steroidogenesis in mast cells, regulating multiple inflammation-associated gene pathways. This study also revealed the uncoupling of transcription and translation, genome and proteome-wide differences of bone marrow and peritoneal cavity-derived mast cells, and striking gender-specific differences at the protein level. This study not only unveils previously unknown phenomena in mast cell biology but also provides a valuable resource for understanding the comparative aspects of mast cells across different origins, genders, and functional states. Our findings have potential translational implications for controlling mast cell-associated pathologies and open new avenues for therapeutic interventions.

## RESULTS

### Mast cells express Cyp11a1 and produce the steroid pregnenolone

The *de novo* steroidogenic potential of immune cells under homeostatic conditions remains unexplored. We investigated Cyp11a1 expression in peripheral tissues and organs using Cyp11a1-mCherry reporter mice (Figure 1A). We found that tissue-resident mast cells in all examined tissues - skin, peritoneal lavage, intestine, lung, adrenal gland, and ovary - express Cyp11a1 to varying degrees, with the highest expression in peritoneal lavage and lowest in the lung (Figure 1B, C). To comprehensively assess Cyp11a1 expression in mast cells across diverse tissues, we analysed publicly available single-cell transcriptomics data (Tauber et al., 2023). We visualised *Cyp11a1* expression levels and the percentage of *Cyp11a1*-expressing mast cells in various mouse organs, including adult stomach, gut mucosa, heart, mammary gland during pregnancy, neonatal muscle, skin, and uterus, peritoneal cavity, and small intestine (Supplementary Figure S1A, B). Our analysis revealed that Cyp11a1 expression is a conserved feature of tissue-resident mast cells across diverse organ systems. All examined mast cell populations expressed Cyp11a1, though with notable variations in both the proportion of expressing cells and expression levels between different tissues (Figure 1B, C; Supplementary Figure S1A, B). This finding suggests that Cyp11a1 expression and *de novo* steroidogenesis may be a fundamental and widespread characteristic of mast cell biology, with potential tissue-specific modulation.

Informed by these observations, we differentiated bone marrow-derived mast cells (BMMCs) and peritoneal lavage-derived mast cells (PMCs) *in vitro* (Figure 1D) and as expected, we found they express Cyp11a1 (Supplementary Figure S1C). To test whether Cyp11a1 expression is functional, we differentiated mast cells from Cyp11a1-sufficient and -deficient precursors (Figure 1E, Supplementary Figure S1D) and analysed culture supernatants by mass spectrometry for steroid profiling (Figure 1F). *Cyp11a1* deletion did not dramatically inhibit mast cell differentiation, but we observed a statistically significant reduction in the number of differentiated mast cells, as defined by cKit and FcεRI expression (Figure 1E). Importantly, mature mast cells produced significant quantities of pregnenolone and detectable levels of progesterone (Figure 1F). Genetic deletion of *Cyp11a1* in mast cells abolished both steroids, confirming active catalysis of cholesterol to pregnenolone by Cyp11a1 (Figure 1F). Steroid profiling revealed that mast cells produce detectable levels of other steroids beyond pregnenolone and progesterone, but their production is not dependent on mast cell-intrinsic Cyp11a1 activity. The complete steroid profiling resource can be accessed in the Supplementary Table S1. This suggests that mast cells may possess additional steroid conversion pathways independent of Cyp11a1 activity, which warrants further investigation.

These findings collectively demonstrate that mast cells express functional Cyp11a1 and possess the capacity for *de novo* steroidogenesis, representing a previously unrecognized aspect of mast cell biology. The expression of Cyp11a1 across various tissue-resident mast cells suggests that this primitive form of steroidogenesis may play a crucial role in mast cell function, particularly in the context of immune and tissue homeostasis.

### Transcriptional remodelling during mast cell regranulation reveals subset-specific programmes and steroidogenic priming

To elucidate the global gene expression patterns and their differences during maturation and regranulation, we performed comprehensive transcriptome analysis on bone marrow-derived mast cells (BMMCs) and peritoneal mast cells (PMCs) in both mature and regranulating states (Figure 2A). We defined regranulating mast cells as those harvested 8 hours after IgE-mediated degranulation, a time point chosen based on previous studies showing the initiation of transcriptional and translational changes during this period (Burwen, 1982; Dvorak et al., 1988; Iskarpatyoti et al., 2022; Jayapal et al., 2006; Kuehn et al., 2010; Xiang et al., 2001).

**Figure 2.**
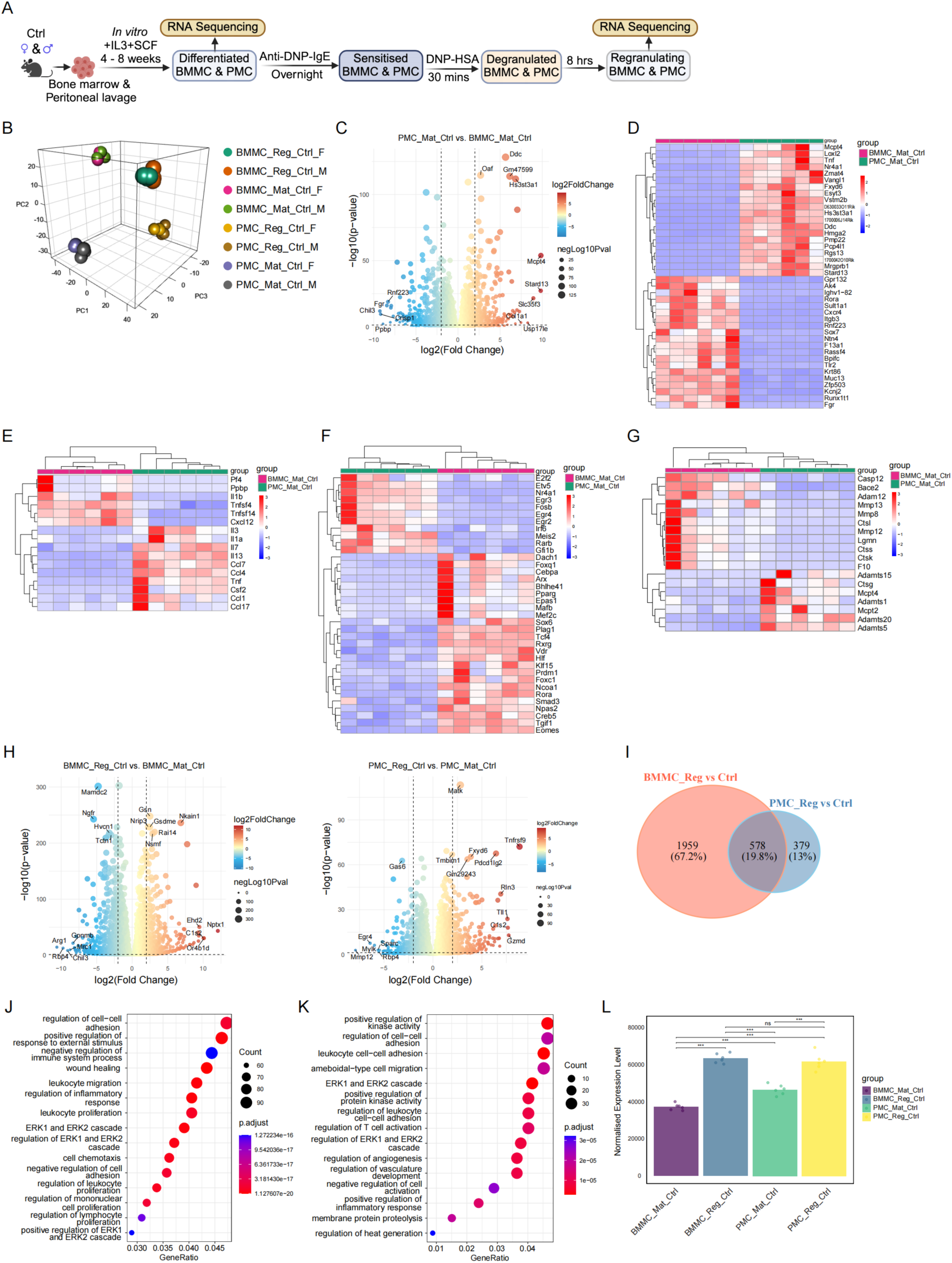
Genome-wide transcriptional comparison of mature and regranulating mast cells in bone marrow derived and peritoneal mast cells (BMMC and PMC). **A.** Schematic representation of the experimental plan. Bone marrow or peritoneal cavity-derived mast cells (BMMC and PMC) were *in vitro* differentiated to obtain mature mast cells. Mature mast cells were sensitised by anti-DNP IgE antibody and activated by exposing them to DNP-HSA. Eight hours after degranulation, cells were considered as regranulating mast cells. Bulk RNA sequencing (RNAseq) of mature and regranulating mast cells (BMMC or PMC) was performed and further analysed bioinformatically. **B.** Principal component analysis (PCA) of RNAseq data. Reg: Regranulating cells (8 hours after degranulation), Mat: Mature, Ctrl: Control (*Cyp11a1*^fl/fl^), M: Male, F: Female. **C.** Volcano plot showing differentially expressed genes in mature BMMC and PMC. Each dot represents an individual gene. Red dots indicate significantly upregulated genes), blue dots indicate significantly downregulated genes, and grey dots represent genes that did not meet the significance threshold. Selected differentially expressed genes are labelled. **D.** Heatmap of top 20 differentially expressed, upregulated or downregulated, genes as revealed in C. **E.** Heatmap of differentially expressed cytokines in mature BMMC and PMC. **F.** Heatmap of differentially expressed transcription factors in mature BMMC and PMC. **G.** Heatmap of differentially expressed proteases in mature BMMC and PMC. **H.** Volcano plot representation of differentially expressed genes during regranulation of mast cells compared to their mature state (BMMC, left panel, and PMC, right panel). Each dot represents an individual gene. Red dots indicate significantly upregulated genes, blue dots indicate significantly downregulated genes and grey dots represent genes that did not meet the significance threshold. Selected differentially expressed genes are labelled. **I.** Venn diagram depicting the similarities and differences in differentially expressed genes (as observed in H) during BMMC and PMC regranulation. **J.** Gene Ontology (GO) enrichment on the differentially expressed genes during BMMC regranulation (as observed in H, left) **K.** Gene Ontology (GO) enrichment analysis on the differentially expressed genes during PMC regranulation (as observed in H, right) **L.** Graphical representation of Cyp11a1 expression in PMC and BMMC. Cyp11a1 increases during regranulating phase, both in PMC and BMMC. P-values are calculated using pairwise t-tests between groups for the expression data with the compare_means function from the ggpubr package (version 0.6.0). The p-values are converted into significance stars (*** for p ≤ 0.001, ** for p ≤ 0.01, * for p ≤ 0.05, and ns for p > 0.05) for annotation on the plot.

Principal component analysis (PCA) revealed distinct transcriptional profiles between BMMCs and PMCs, with significant changes occurring during the regranulation phase (Figure 2B). Interestingly, while there were clear differences between cell types and activation states, transcriptomic differences between male and female cohorts were minimal, allowing us to combine male and female samples for subsequent analyses.

Differential gene expression analysis uncovered substantial differences between mature BMMCs and PMCs (Figure 2C, Supplementary Table S2). Among the top 20 differentially expressed genes, markers such as *Mcpt4* (highly expressed in PMCs) and *Cxcr4* (enriched in BMMCs) were prominent, validating our experimental approach. Additionally, novel genes such as *Tnf*, *Nr4a1*, *Stard13*, and *Ddc* were highly expressed in PMCs, while *Gpr132*, *Rora*, *Sult1a1*, *Fgr*, and *Chil3* were enriched in BMMCs, highlighting tissue-specific adaptations (Figure 2D). Further categorization of differentially expressed genes revealed variations in cytokines, transcription factors, and proteases between the two cell types. For instance, BMMCs showed higher expression of pro-inflammatory cytokines *Pf4*, *Ppbp*, *Il1b*, *Tnfsf14* and *Cxcl12* compared to PMCs, which expressed higher levels of cytokines such as *Il3*, *Il7*, *Il13*, and chemokines like *Ccl7* and *Ccl17* (Figure 2E). Transcription factors such as *Nr4a1* and *Rora* were differentially regulated between the two populations (Figure 2F), while proteases such as *Mcpt4* (PMC-specific) and *Caasp12* (BMMC-specific) further underscored their functional specialization (Figure 2G). These differences likely contribute to the functional specialization of mast cells in different tissue environments.

The most dramatic transcriptomic changes were observed when comparing regranulating mast cells to their mature counterparts (Figure 2H, Supplementary Table S3 for the left panel, Supplementary Table S4 for the right panel). This analysis identified both previously known degranulation-associated genes and novel candidates, significantly expanding our understanding of the molecular events during this critical phase. Venn diagram analysis revealed 578 common differentially expressed genes between regranulating BMMCs and PMCs, along with 1959 BMMC-specific and 379 PMC-specific genes (Figure 2I). This suggests both shared and distinct regranulation programmes in these two mast cell populations. To elucidate the functional implications of transcriptional changes during regranulation, we conducted Gene Ontology (GO) enrichment analysis on differentially expressed genes in BMMCs and PMCs (Figure 2J, K). Both cell types displayed significant enrichment in biological processes related to cell-cell adhesion, negative regulation of immune responses, and wound healing, highlighting shared pathways involved in their regranulation dynamics and regranulating phase-specific function. Notably, the ERK1/ERK2 signalling cascade emerged as a prominently enriched pathway in both BMMCs and PMCs, underscoring its pivotal role in orchestrating the regranulation process. However, distinct transcriptional profiles were also observed between the two cell types. For instance, BMMCs exhibited stronger enrichment in processes such as leukocyte migration and inflammatory response regulation, while PMCs showed unique enrichment in pathways like positive regulation of kinase activity and membrane protein proteolysis. These findings underscore both shared and cell type-specific mechanisms underlying mast cell regranulation.

Notably, we observed that *Cyp11a1* expression increased during the regranulation phase in both PMCs and BMMCs (Figure 2L), indicating a potential role for steroidogenesis in this process. This finding connects our transcriptomic data with the steroidogenic capacity of mast cells described earlier.

Collectively, this genome-wide expression analysis provides a comprehensive view of the transcriptional landscape in mature and regranulating mast cells. It not only confirms known aspects of mast cell biology but also uncovers novel genes and pathways involved in mast cell function and regulation. The identification of distinct transcriptional programmes in BMMCs and PMCs, as well as the shared and unique responses during regranulation, offers valuable insights into the molecular basis of mast cell heterogeneity and plasticity. These findings lay the groundwork for future studies aimed at understanding and potentially modulating mast cell functions in various physiological and pathological contexts.

### *De novo* steroidogenesis orchestrates mast cell development and functional recovery through distinct transcriptional programmes

To delineate the cell-intrinsic role of steroidogenesis in mast cell biology, we performed transcriptome analysis of control and *Cyp11a1*-deficient BMMCs and PMCs (Figure 3A). Principal component analysis revealed distinct transcriptional signatures between control and *Cyp11a1*-deficient cells, with the most pronounced differences emerging during the regranulation phase (Figure 3B). Differential gene expression analysis uncovered widespread transcriptional alterations by *Cyp11a1* deletion, with 321 differentially expressed genes in BMMCs (199 upregulated, 122 downregulated) and 664 in PMCs (221 upregulated, 443 downregulated) (Figures 3C-D, Supplementary Tables S5-S6). The marked predominance of downregulated genes in PMCs suggests that Cyp11a1-dependent steroidogenesis primarily functions as a positive regulator of gene expression in tissue-resident mast cells.

**Figure 3.**
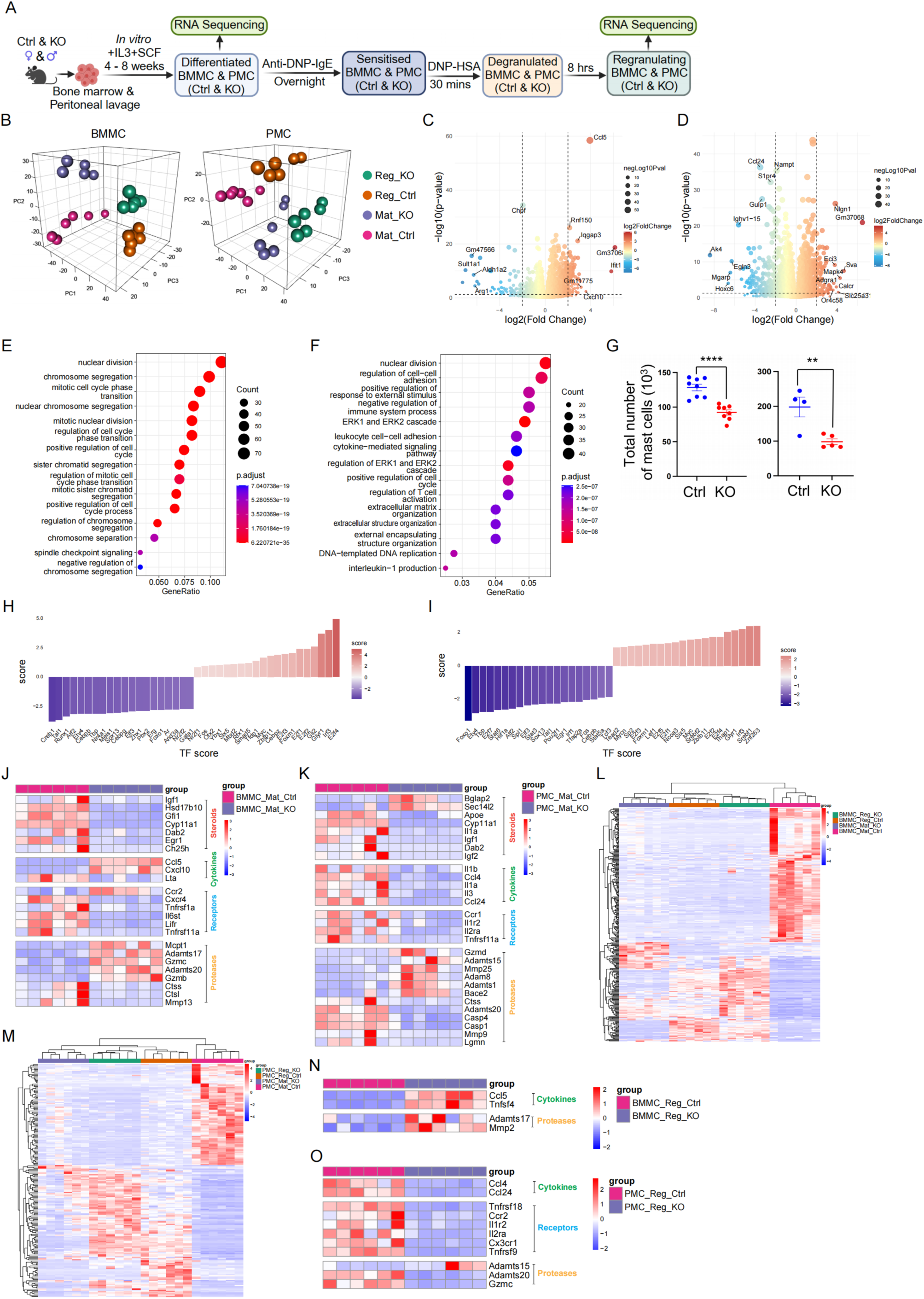
Genome-wide gene regulatory effect of *Cyp11a1* deletion on mast cell differentiation and regranulation. **A.** Schematic representation of the experimental plan. Control (*Cyp11a1*^fl/fl^) or *Cyp11a1* knockout (KO) BMMC and PMC were *in vitro* differentiated to obtain mature mast cells. Mature mast cells were sensitised by anti-DNP IgE antibody and activated by exposing them to DNP-HSA. Eight hours after degranulation cells were considered and regranulating mast cells. Bulk RNA sequencing (RNAseq) of mature and regranulating mast cells was performed, and further analysed bioinformatically. **B.** Principal component analysis (PCA) of the transcriptomes. Reg: Regranulating, Ctrl: Control *Cyp11a1* ^fl/fl^, KO: *Cyp11a1* knockout mast cells. **C.** Volcano plot depicting differentially expressed genes in control and *Cyp11a1* knockout BMMC. **D.** Volcano plot of differentially expressed genes in control and *Cyp11a1* knockout PMC. **E.** Gene set enrichment (GO Gene Ontology) analysis shows the biological processes in BMMC affected by the *Cyp11a1* knockout. **F.** Gene set enrichment (GO Gene Ontology) analysis shows the biological processes in PMC affected by the *Cyp11a1* knockout. **G.** An equal number of mast cells were plated for BMMC or PMC differentiation. *Cyp11a1* KO mast cell numbers were reduced compared to control mast cells. Representative of four independent experiments. p-value was calculated using unpaired two-tailed t-test. **H.** Bar graph representation of the differentially regulated transcription factors (TF) in BMMC because of *Cyp11a1* deletion. **I.** Bar graph representation of the differentially regulated transcription factors (TF) in PMC because of *Cyp11a1* deletion. **J.** Heatmap representation of the differentially regulated steroid pathway-associated genes, cytokines, receptors and proteases in BMMC because of *Cyp11a1* deletion. **K.** Heatmap representation of the differentially regulated steroid-associated pathway genes, cytokines, receptors and proteases in PMC because of *Cyp11a1* deletion. **L.** Heatmap showing differentially regulated "regranulation-associated regulatory genes" in BMMC following *Cyp11a1* deletion. Genes differentially expressed during regranulation, as identified in Figure 2H (left panel), were classified as "regranulation-associated regulatory genes." The heatmap visualizes the impact of *Cyp11a1* knockout on the expression of these genes. Heatmap representation of the differentially regulated “regranulation-associated regulatory genes”, as found in Figure 2H (right panel) in PMC because of *Cyp11a1* deletion. **M.** Heatmap representation of the differentially regulated cytokines and proteases among the differentially regulated “regranulation-associated regulatory genes” in BMMC (as revealed in L) because of *Cyp11a1* deletion. **N.** Graphical representation of the differentially regulated cytokines, receptors and proteases among the differentially regulated “regranulation-associated regulatory genes” in PMC (as revealed in M) because of *Cyp11a1* deletion.

Gene set enrichment analysis revealed that Cyp11a1-regulated genes primarily control cell division, cell cycle, and immune response-associated processes in both BMMCs and PMCs (Figure 3E-F). This indicates that mast cell steroidogenesis plays a crucial role in regulating cell proliferation during maturation and potentially effector functions. In PMCs, the enrichment pattern demonstrated a broader impact on cellular functions (Figure 3F). The analysis revealed significant enrichment in pathways related to nuclear division, cell-cell adhesion, immune system processes, and ERK1/2 cascade regulation. Notably, several immune-related processes were prominently affected, including leukocyte cell-cell adhesion, cytokine-mediated signalling, and T cell activation regulation, indicating that *Cyp11a1* deletion substantially impacts the immune regulatory functions of tissue-resident mast cells. The functional significance of Cyp11a1 in mast cell development was further demonstrated by cellular enumeration experiments (Figure 3G). When equal numbers of cells were initially plated, both BMMC and PMC populations showed significantly reduced cell numbers in *Cyp11a1*-deficient conditions compared to controls. This reduction in cell numbers aligns with the GO analysis findings, particularly the enrichment of cell cycle-related pathways, and suggests that *Cyp11a1*-dependent steroidogenesis is essential for optimal mast cell proliferation, development and survival.

*Cyp11a1* deletion triggered distinct alterations in transcription factor networks between BMMCs and PMCs (Figures 3H-I), with differential regulation of key developmental regulators including *Creb1, Tal1,* and *Runx1* in BMMCs, and *Foxo1, Etv4,* and *Tbp* in PMCs. Interestingly, Irf3 has been found to upregulated in both BMMC and PMC. IRF3 is known to enhance mast cell degranulation and histamine release during IgE-mediated allergic reactions. These changes were accompanied by cell type-specific modulation of steroid signalling components, cytokines, receptors, and proteases (Figures 3J-K). In BMMCs, Cyp11a1 deficiency led to downregulation of steroid signalling genes (*Igf1, Hsd17b10, Gfi1*) while differentially affecting inflammatory mediators and proteases. PMCs exhibited a distinct response pattern, characterized by consistent downregulation of cytokines (*Il1b, Ccl4, Il3*) and cell type-specific alterations in protease expression. These findings highlight the cell type-specific effects of *Cyp11a1* deletion on mast cell function, suggesting that the abrogation of mast cell-mediated steroidogenesis differentially impacts the transcriptional programmes governing inflammation, cell signalling, and tissue remodelling in BMMCs and PMCs.

A particularly striking finding emerged from our analysis of regranulation-associated genes - those specifically modulated during the recovery phase after degranulation as observed in Figure 2H. The heatmap analysis revealed that *Cyp11a1* deletion substantially altered the expression patterns of these regranulation-specific genes in both BMMCs and PMCs (Figures 3L-M, Supplementary Table S7, S8). In BMMCs, Cyp11a1 deficiency led to a distinct reorganization of the regranulation-associated transcriptional programme, with clear clustering patterns distinguishing control from knockout cells. A striking observation emerged: *Cyp11a1* deletion pre-emptively altered the expression patterns of regranulation-associated genes, mirroring their expected regulatory changes during the normal regranulation phase. These genes then displayed further differential expression during the actual regranulation process, suggesting that Cyp11a1-mediated steroidogenesis primes mast cells for appropriate functional recovery after degranulation. This pattern is clearly visualized in the heatmaps (Figures 3L-M). A complete comparative heatmap of all differentially expressed genes has been shown in Supplementary Figure S3A-B (Supplementary Table S9,10). Detailed examination revealed significant dysregulation of key mediators, particularly affecting *Ccl5* and *Tnfsf4* expression, along with the proteases *Adamts17* and *Mmp2* (Figure 3N), suggesting disruption of both inflammatory mediator production and tissue remodelling capabilities during the recovery phase. In PMCs, the impact of Cyp11a1 deficiency on regranulation-associated genes was even more pronounced, with substantial dysregulation of multiple functional categories (Figure 3O): cytokines (*Ccl4, Ccl24*), receptors (*Il1r2, Il2ra Tnfrsf18/192, Cx3cr1*), and proteases (*Adamts15/20, Gzmc*). This comprehensive alteration of the regranulation-associated transcriptional programme suggests that mast cell-derived steroids serve as master regulators of the recovery phase, coordinating both inflammatory resolution and the restoration of cellular functionality.

Collectively, these results establish mast cell steroidogenesis as a central regulator of differentiation, survival, and functional responses, with particular importance in the regranulation phase. The identification of these Cyp11a1-dependent transcriptional programmes provides new insights into the molecular mechanisms governing mast cell functional restoration and suggests potential therapeutic targets for modulating mast cell-mediated inflammatory responses.

### Proteome-wide comparative proteomics reveals sex-specific and state-dependent protein repertoire in mast cells

To complement our transcriptomic findings and gain deeper insights into the functional changes occurring during mast cell regranulation, we performed quantitative proteomics on mature and regranulating BMMCs and PMCs (Figure 4A). Intriguingly, while our transcriptomic analysis showed minimal gender-specific differences, principal component analysis of the proteomics data revealed significant sexual dimorphism in protein expression profiles (Figure 4B). This observation underscores the importance of considering sex as a biological variable in mast cell research and highlights the potential limitations of transcriptome-only studies.

**Figure 4.**
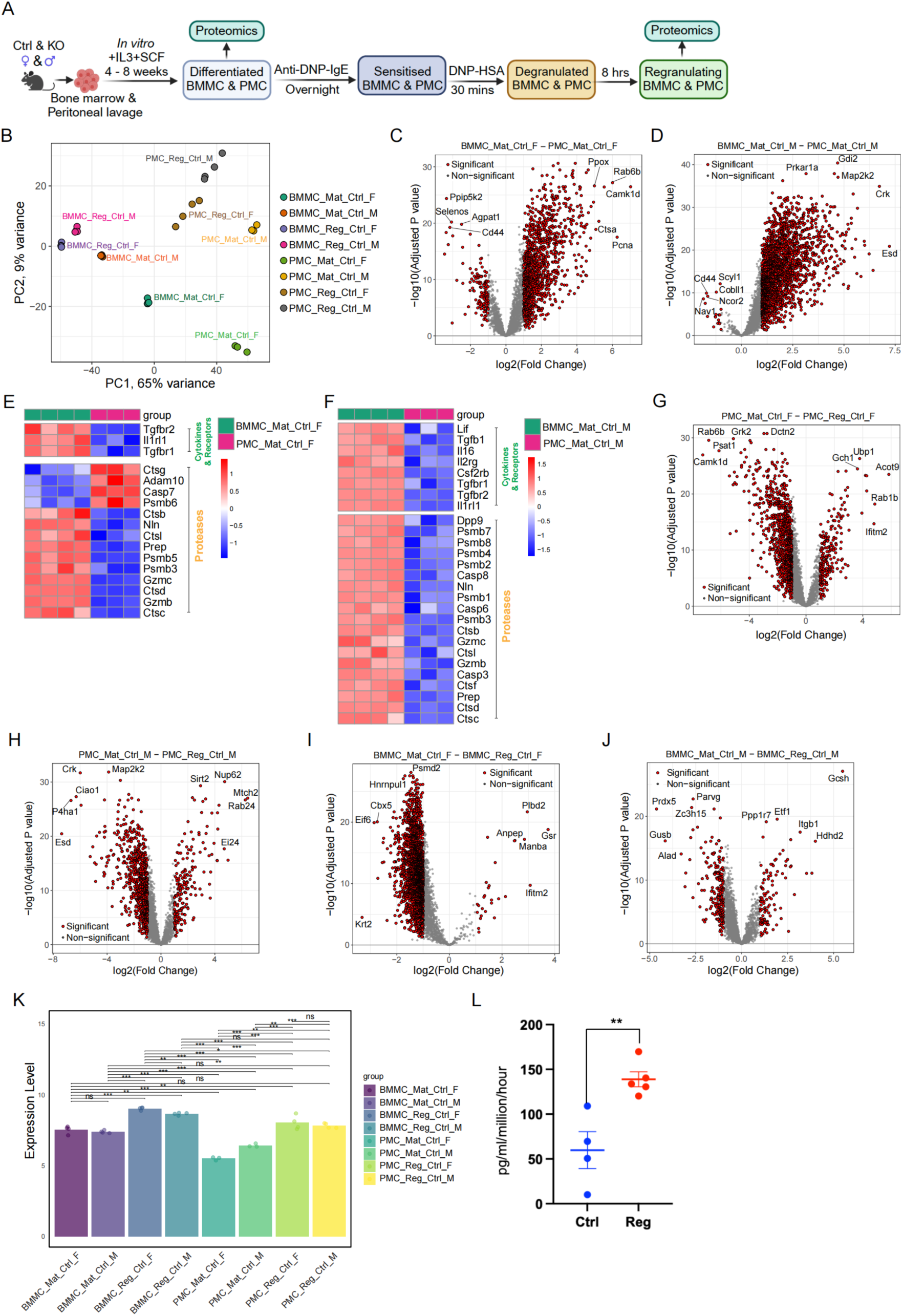
Proteome-wide comparative analysis of mature and regranulating mast cells across sex and tissue origin. **A.** Schematic representation of the experimental plan. Bone marrow or peritoneal cavity-derived mast cells (BMMC and PMC) were *in vitro* differentiated to obtain mature mast cells. Mature mast cells were sensitised by anti-DNP IgE antibody and activated by exposing them to DNP-HSA. Eight hours after degranulation, cells were considered and regranulating mast cells. Mature and regranulating mast cells obtained from BMMC or PMC were used in liquid chromatography, followed by tandem mass-spectrometry-based proteomics detection and subsequent bioinformatics analyses. **B.** Principal component analysis (PCA) of proteomics data. M: Male, F: Female, Mat: Mature, Reg: Regranulating cells, Ctrl: Control *Cyp11a1*^fl/fl^ **C.** Volcano plot showing differential expression of proteins in female mature BMMC and PMC. **D.** Volcano plot depicting differential protein expression in male mature BMMC and PMC. **E.** Volcano plot showing differentially expressed cytokines, repeptors and proteases that are differentially expressed in female BMMC and PMC. **F.** Volcano plot of differentially expressed cytokines, repeptors and proteases that are differentially expressed in male BMMC and PMC. **G.** Volcano plot depicting differential expression of proteins in female mature PMC and regranulating PMC. **H.** Volcano plot showing differentially expressed proteins in male mature PMC and regranulating PMC. **I.** Volcano plot showing differential expression of proteins in female mature and regranulating BMMC. **J.** Volcano plot showing differential expression of proteins in male mature and regranulating BMMC. **K.** The bar graph shows that Cyp11a1 protein level increased during the regranulating phase both in BMMC and PMC. P-values are calculated using pairwise t-tests between groups for the expression data with the compare_means function from the ggpubr package (version 0.6.0). The p-values are converted into significance stars (*** for p ≤ 0.001, ** for p ≤ 0.01, * for p ≤ 0.05, and ns for p > 0.05) for annotation on the plot. **L.** An equal number of mature and regranulating PMCs were cultured parallelly & simultaneously. Cell culture supernatant was analysed by ELISA for pregnenolone production and synthesis rate. Representative of three independent experiments. p-value was calculated using unpaired two-tailed t-test

Given these gender-specific differences, we analysed male and female samples separately. In female mice, we identified 1228 differentially expressed proteins between mature BMMCs and PMCs, with 128 proteins showing higher expression in BMMCs and 1100 proteins more abundant in PMCs (Figure 4C, Supplementary Table S11). Similarly, in male mice, we found 17 proteins highly expressed in BMMCs compared to PMCs, and 2153 proteins more abundant in PMCs (Figure 4D, Supplementary Table S12). Detailed analysis of functionally relevant protein categories revealed distinct patterns between BMMC and PMC populations. In female mice, several key cytokines, receptors, and proteases showed tissue-specific expression patterns (Figure 4E). Notable differences included elevated levels of Il1r1, Tgfbr1/2, and specific proteases like Gzmb/c and Ctsb/c/l in BMMCs, while PMCs exhibited higher expression of distinct protease profiles, including Ctsg and Adam10. Male mice demonstrated similar tissue-specific protein signatures, with differential expression of immune mediators and proteolytic enzymes (Figure 4F), though the specific protein sets differed from females, highlighting gender-specific specialization of mast cell populations.

We next compared the proteomes of mature and regranulating mast cells for both male and female BMMCs and PMCs (Figure 4G-J). This analysis uncovered substantial proteomic remodelling during the regranulation phase, with 1083 significantly differentially expressed proteins, of which 876 proteins were upregulated and 207 proteins downregulated in female PMCs (Figure 4G, Supplementary Figure S13). Similar patterns were observed in male PMCs (Figure 4H, Supplementary Figure S14) and BMMCs of both sexes (Figure 4I-J, Supplementary Figure S15, S16). Interestingly, during regranulation, mast cells upregulate more proteins than they downregulate (Figure 4G-J, Supplementary Table S13-S16), which contrasts with the predominantly downregulated mRNA expression observed in our transcriptomics analysis (Figure 2H, Supplementary Table S3, S4), suggesting temporal uncoupling of transcription and translation.

Of particular interest, we found that Cyp11a1 protein levels increased significantly during the regranulation phase in both BMMCs and PMCs (Figure 4K). This increase in Cyp11a1 protein correlated with enhanced pregnenolone production in regranulating mast cells, as measured by quantitative ELISA (Figure 4L). These findings suggest that steroidogenesis may play a crucial role in the regranulation process and the restoration of mast cell function following activation.

Collectively, our proteomics data provide a comprehensive view of the dynamic changes in protein expression that occur during mast cell regranulation, revealing both tissue-specific and sex-dependent protein signatures. The observed sexual dimorphism and the discrepancies between transcriptomic and proteomic profiles highlight the complexity of mast cell biology and emphasise the importance of integrative multi-omics approaches. Moreover, the upregulation of Cyp11a1 and increased steroid production during regranulation further support a central role for de novo steroidogenesis in mast cell function and recovery following activation.

### *De novo* steroidogenesis regulates sex-specific proteomic landscape of mast cells: From homeostasis to regranulation

To elucidate the impact of mast cell steroidogenesis on protein expression, we performed quantitative proteomics on control (*Cyp11a1*^fl/fl^) and *Cyp11a1* knockout mature and regranulating BMMCs and PMCs (Figure 5A). This analysis revealed substantial alterations in protein expression patterns due to *Cyp11a1* deletion in both mature and regranulating states of BMMCs and PMCs, with distinct changes observed between male and female samples (Figure 5B-O). In mature female BMMCs, *Cyp11a1* deletion resulted in the differential expression of 178 proteins (80 upregulated and 98 downregulated (Figure 5B, Supplementary Table S17). Male BMMCs showed more modest changes, with 30 differentially expressed proteins (14 upregulated, 16 downregulated; Figure 5C, Supplementary Table S18). PMCs exhibited more extensive alterations, with 329 and 136 proteins differentially expressed in female and male samples, respectively (Figure 5D-E, Supplementary Figure S119, S20).

**Figure 5.**
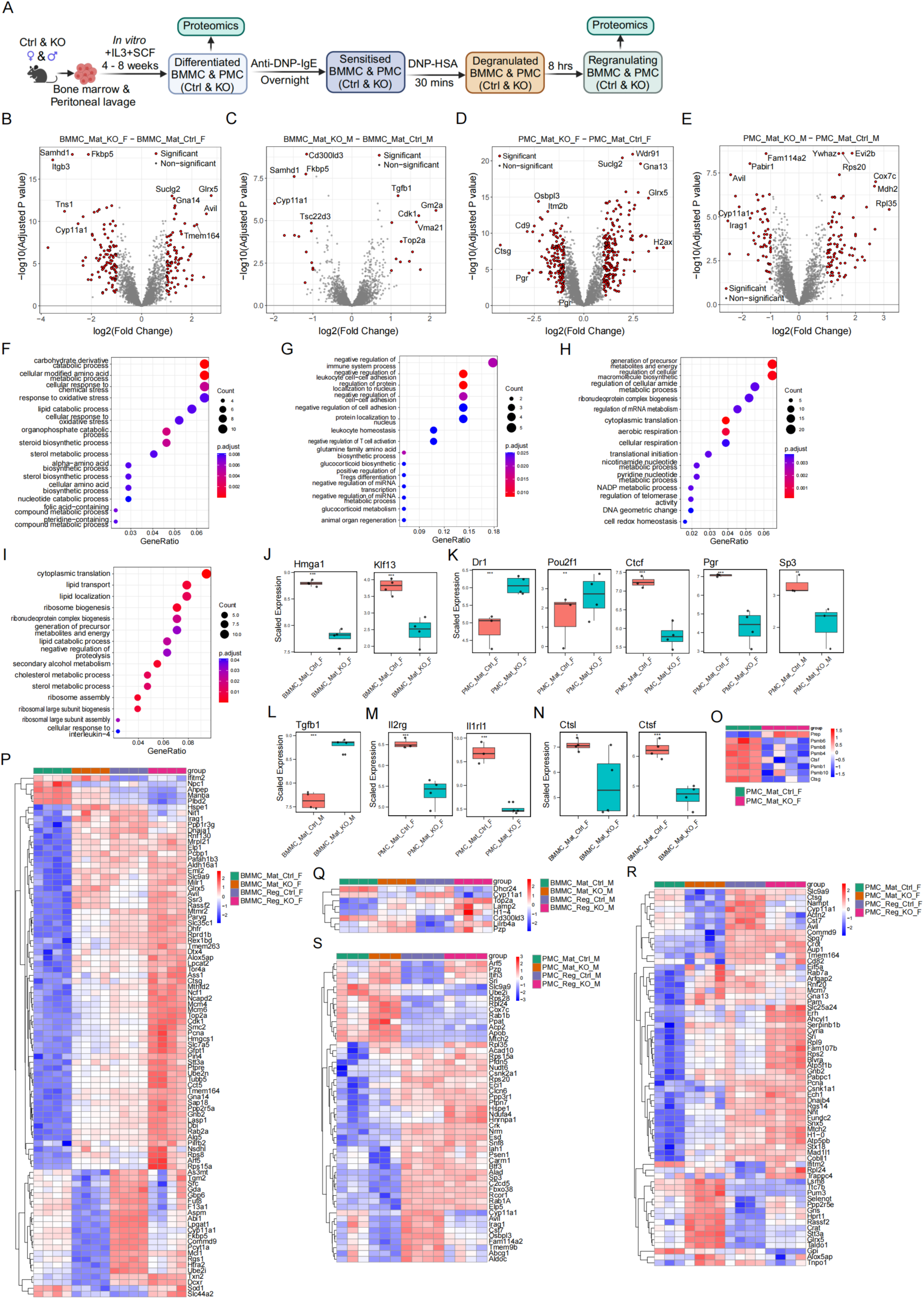
Mast cell steroidogenesis regulates cell-intrinsic translational-level regulatory circuits of mast cell maturation and function. **A.** Schematic representation of the experimental plan. Control (Cyp11a1^fl/fl^) or *Cyp11a1* knockout (KO) BMMC and PMC were *in vitro* differentiated to obtain mature mast cells. Mature mast cells were sensitised by anti-DNP IgE antibody and activated by exposing them to DNP-HSA. Eight hours after degranulation, cells were considered and regranulating mast cells. Liquid chromatography tandem mass spectrometry based proteomics of KO or control mature and regranulating BMMC or PMC was performed, and further bioinformatics analyses were used. **B-E.** Volcano plots show proteome-wide differentially expressed proteins due to *Cyp11a1* deletion in mature female BMMC (B), male BMMC (C), female PMC (D) and male PMC (E). **F-I.** GO enrichment analysis in differentially expressed genes upon *Cyp11a1* knockout in mature female BMMC (F), male BMMC (G), female PMC (H) and male PMC (I) **J.** The graphical representation shows transcription factors’ expression affected by *Cyp11a1* knockout in female BMMC **K.** A graphic representation shows transcription factors’ expression affected by *Cyp11a1* knockout in mature female (first four panels) and male (last panel, Sp3) PMC. **L.** The graphical representation shows cytokines’ expression affected by *Cyp11a1* knockout in mature female (top) and male (bottom) BMMC. **M.** The graphical representation shows cytokine and cytokine receptor expression affected by *Cyp11a1* knockout in mature female PMC. **N.** Graphical representation shows proteases’ expression affected by *Cyp11a1* knockout in mature female BMMC. **O.** Heatmap shows proteases’ expression affected by *Cyp11a1* knockout in mature female PMC.

Gene Ontology (GO) enrichment analysis of the differentially expressed proteins revealed significant alterations in biological processes and molecular functions in both BMMCs and PMCs (Figure 5F-I). In BMMCs, Cyp11a1 activity primarily influenced cellular metabolism and immune response pathways, while in PMCs, cellular metabolism and protein synthesis-associated factors were affected.

Further analysis of specific protein categories revealed that *Cyp11a1* deletion affected the expression of several transcription factors in both BMMCs and PMCs (Figure 5J-K). Notably, in female BMMCs, factors such as Klf13 and Hmga1 were significantly downregulated (Figure 5J), while in female PMCs, down regulator of transcription (Dr1), Pou2f1, Ctcf, and progesterone receptor (Pgr) were downregulated (Figure 5K, first 4 panels). Sp3 was downregulated in male PMC (Figure 5K, last panel). These changes in transcriptional regulators likely contribute to the broad proteomic alterations observed and may explain the extensive transcriptomic changes reported earlier. The expression of cytokines and cytokine receptors was also modulated by *Cyp11a1* deletion (Figure 5L-M). In BMMCs, Tgfb1 showed significant upregulation (Figure 5J), while in PMCs, Il2rg and Il1rl1 were differentially expressed (Figure 5M). These alterations in cytokine and their receptor expression profiles could have substantial implications for mast cell-mediated immune responses and intercellular communication. Interestingly, we observed changes in the expression of proteases upon *Cyp11a1* deletion (Figure 5N, O). In female BMMCs, cathepsins such as Ctsl and Ctsf (Figure 5N) showed differential expression, while in female PMCs, apart from cathepsin, a broader range of proteases was affected (Figure 5O).

Intriguingly, the impact of *Cyp11a1* deletion on protein expression was interesting in regranulating mast cells. In female BMMCs, the effect was more severe than male BMMC (Figure 5P-Q). We observed four patterns of protein expression changes (Figure 5P): (1) Protein factors that are typically upregulated during regranulation were already upregulated in *Cyp11a1*-deficient BMMCs, and these proteins were further upregulated during regranulation phage. (2) Proteins that are upregulated during regranulation were downregulated upon *Cyp11a1* knockout and did not respond as of wild-type cells in elevating the protein level. (3) The proteins that are typically downregulated during regranulation, were already downregulated upon *Cyp11a1* knockout cells and further downregulated upon regranulation phase. (4) Proteins, at least two proteins, Sod1 and Slc44a2, which are downregulated in regranulation phase, however, upregulated in *Cyp11a1* knockout cells. In male BMMC, we observed a similar pattern but with only a few proteins (Figure 5Q). PMCs exhibited similar trends and protein expression patterns but with varying degrees because of *Cyp11a1* deletion (Figure 5R-S). Importantly, we observed that male PMCs are more affected (Figure 5S) than male BMMC (Figure 5Q). This enhanced effect of *Cyp11a1* knockout during regranulation suggests that mast cell steroidogenesis plays a crucial role in shaping the protein landscape during functional recovery after degranulation.

In conclusion, our proteomic analysis reveals that mast cell steroidogenesis profoundly influences the cellular protein landscape, regulating key factors involved in transcriptional control, cytokine signalling, and proteolytic functions. The heightened impact observed during regranulation underscores the critical role of this pathway in mast cell functional recovery and homeostasis.

### Global uncoupling of transcriptional and translational programmes in mast cell function

Our integrative analysis of transcriptomes and proteomes revealed a striking disconnect between mRNA and protein expression patterns across mast cell states and genotypes. Comparative analysis of differentially expressed genes (DEGs) and proteins (DEPs) showed remarkably limited overlap across all conditions examined (Figures 6A-D). In BMMCs, only 28 molecules (0.922%) were identified as both DEGs and DEPs when comparing regranulating to mature states, despite the presence of 2,509 DEGs and 500 DEPs (Figure 6A). This minimal overlap was even more pronounced in PMCs, where only 6 molecules (0.369%) were common between 951 DEGs and 667 DEPs (Figure 6B). The disconnection between transcriptional and translational programmes persisted in the context of Cyp11a1 deletion. In BMMCs, only 10 molecules (1.5%) overlapped between 619 DEGs and 39 DEPs (Figure 6C). Similarly, in PMCs, just 10 molecules (1.38%) were common between 638 DEGs and 76 DEPs (Figure 6D).

**Figure 6.**
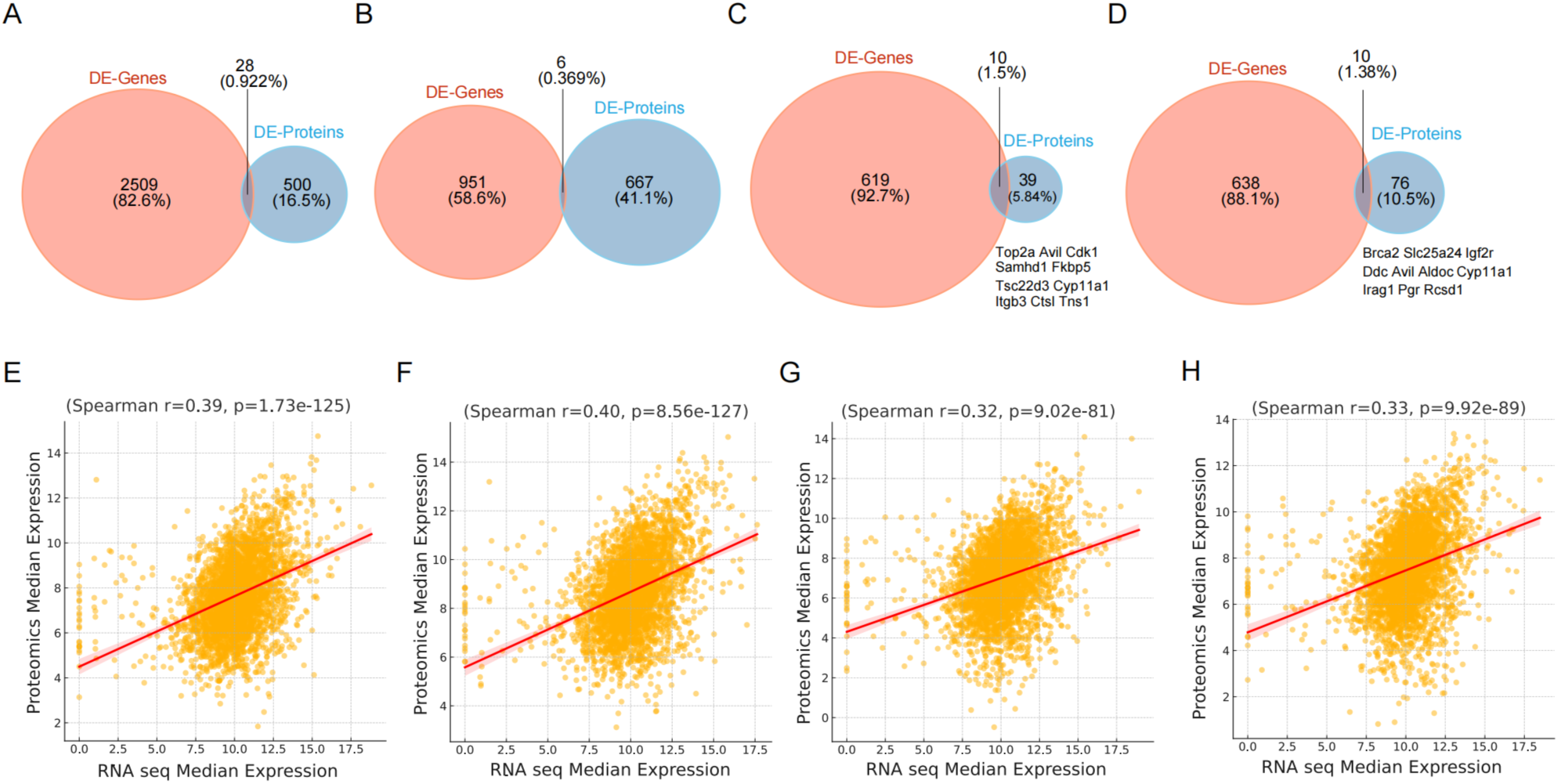
Global uncoupling of transcription and translation in mast cells. **A.** Venn diagram showing overlap of differentially expressed genes (DE-Genes) in the transcriptome and differentially expressed proteins (DE-Proteins) in the mature and regranulating BMMC proteome. **B.** Venn diagram showing overlap of DE-Genes in the transcriptome and DE-Proteins in the proteome of mature and regranulating PMC. **C.** Venn diagram showing overlap of DE-Genes in the transcriptome and DE-Proteins in the proteome of control and *Cyp11a1* KO BMMC. **D.** Venn diagram showing overlap of DE-Genes in the transcriptome and DE-Proteins in the proteome of control and *Cyp11a1* KO PMC. **E-F.** Gene-wise median correlation (Spearman’s rank correlation coefficient) of mRNA and protein expression in mature (E) and regranulating (F) BMMC. **G-H.** Gene-wise median correlation (Spearman’s rank correlation coefficient) of mRNA and protein expression in mature (G) and regranulating (H) PMC.

To quantitatively assess this uncoupling, we performed genome-wide correlation analyses between mRNA and protein expression levels. In BMMCs, Spearman correlation coefficients were notably low in both mature (r = 0.39) and regranulating (r = 0.40) states (Figures 6E-F). PMCs exhibited similarly weak correlations in mature (r = 0.32) and regranulating (r = 0.33) conditions (Figures 6G-H). These consistently low correlation coefficients across all conditions suggest that post-transcriptional regulation serves as a major determinant of protein abundance in mast cells.

This global uncoupling was particularly evident in the regranulation response, where we observed an inverse relationship between transcript and protein regulation patterns. While our transcriptomic analysis revealed a predominance of downregulated genes during regranulation, proteomic data showed a contrasting pattern with more upregulated proteins. This inverse relationship suggests the existence of sophisticated post-transcriptional regulatory mechanisms that fine-tune mast cell functional responses.

These findings reveal an unprecedented layer of complexity in mast cell regulation, where transcriptional and translational programmes operate with substantial independence. This uncoupling may represent a key mechanism allowing mast cells to rapidly modulate their functional states while maintaining long-term transcriptional memory, particularly during the critical process of regranulation.

### *De novo* steroidogenesis functions as a homeostatic regulator of mast cell responses

Our integrative omics analysis identified steroidogenesis as a key regulatory pathway in mast cells, controlling genes essential for immune modulation and cellular homeostasis. To elucidate the functional significance of this pathway, we systematically investigated mast cell responses using complementary in vitro and in vivo approaches.

To test the functional regulatory role of mast cell *de novo* steroidogenesis, we conducted IgE-dependent (using DNP-HSA and anti-DNP IgE) and IgE-independent (compound 48/80 mediated) degranulation assays using BMMC and PMCs derived from Cyp11a1-sufficient (*Cyp11a1*^fl/fl^ control)) or -deficient mice (using *Cyp11a1*^fl/fl^;*Vav1*^Cre^ mice or *Cyp11a1*^fl/fl^;*Cpa3*^Cre^) (Figure 7A). BMMC and PMCs were sensitised by anti-DNP-HSA IgE antibody and activated by DNP-HSA (Figure 7A). Degranulation was measured by β-hexosaminidase assay. We observed a significant increase in degranulation in *Cyp11a1* knockout mast cells (Figure 7B). We observed a similar pattern of results when mast cells were activated in an IgE-independent manner using compound 48/80 (Figure 7C). These results suggest the potential absence of a Cyp11a1-dependent negative regulatory signalling pathway.

**Figure 7.**
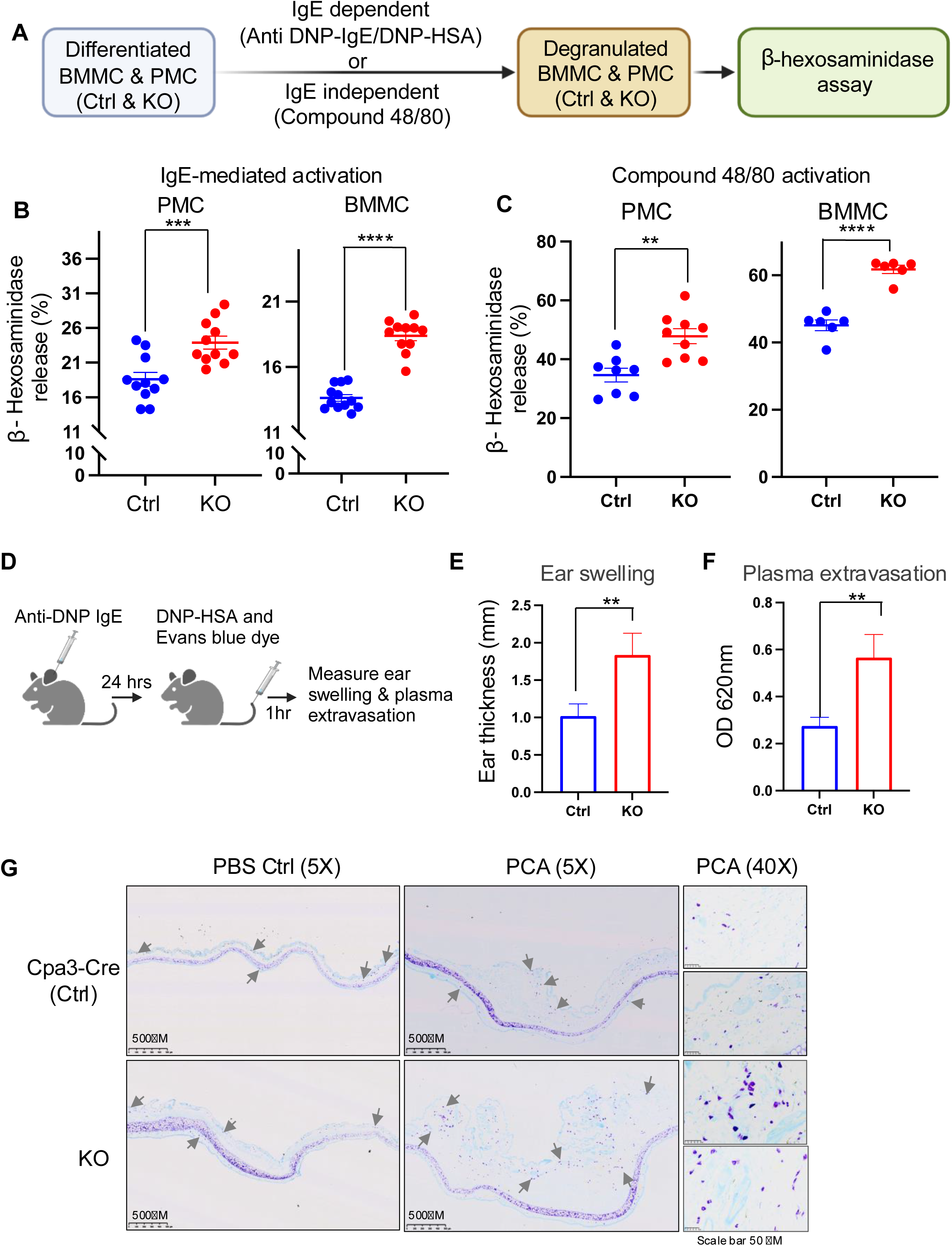
Mast cell steroidogenesis acts as a negative regulatory mechanism to control inflammatory responses. **A.** Schematic of experimental design for assessing mast cell degranulation in vitro. Control (*Cyp11a1*^fl/fl^) and *Cyp11a1*-deficient mast cells were activated via IgE-dependent (anti-DNP IgE/DNP-HSA) or IgE-independent (compound 48/80) pathways. Degranulation was quantified by β-hexosaminidase release assay. **B.** Enhanced IgE-mediated degranulation in *Cyp11a1*-deficient PMCs (left) and BMMCs (right) compared to controls, measured by β-hexosaminidase release. Representative of three independent experiments. p-value was calculated using unpaired two-tailed t-test. **C.** Increased IgE-independent degranulation in *Cyp11a1*-deficient PMCs (left) and BMMCs (right) in response to compound 48/80. Representative of three independent experiments. p-value was calculated using unpaired two-tailed t-test. **D-G.** *Cyp11a1* deletion in mast cells exacerbates passive cutaneous anaphylaxis (PCA). (D) Experimental design: Ears of *Cpa3*^Cre^ control and *Cyp11a1*^fl/fl^;*Cpa3*^Cre^ mice were sensitized with anti-DNP IgE, followed by intravenous injection of DNP-HSA and Evans blue dye. (E) Increased ear swelling and (F) enhanced vascular permeability (Evans blue extravasation) in *Cyp11a1*-deficient mice compared to controls. (G) Representative Toluidine blue staining showing increased degranulating mast cells and tissue inflammation in *Cyp11a1*-deficient mice. A. Representative of three independent experiments. p-value was calculated using unpaired two-tailed t-test

To validate these findings *in vivo*, we conducted passive cutaneous anaphylactic (PCA) reactions in mast cell & basophil-specific *Cyp11a1* knockout mice using *Cyp11a1*^fl/fl^ ;*Cpa3*^Cre^ mice (KO). This *Cpa3*^Cre^ line is the only Cre-driver mouse line to delete mast cell-specific genes in all mast cells (Lilla et al., 2011) across mucosal and connective tissues. Cpa3 is also expressed in basophils. However, the inflammatory phenotypes of the PCA reaction model used here are well-established for mast cell-mediated responses. *Cpa3*^Cre^ deletes *Cyp11a1* in mast cells almost entirely (Supplementary Figure S7A). In these *Cpa3*^Cre^-driven *Cyp11a1* knockout mice, blood basophil density remains unchanged (Supplementary Figure S7B). We did not observe changes in the frequency and localisation of mast cells in any observed tissues, such as the tongue, intestine and ear skin of the KO mice compared to control mice (Supplementary Figure S7C). In PCA experiments, ear resident mast cells were sensitised by anti-DNP IgE injection (Figure 7D). The next day, cutaneous anaphylaxis was induced by intravenous injection of DNP-HSA (along with Evans blue dye to monitor plasma extravasation). We observed markedly enhanced inflammatory responses in Cyp11a1-deficient (KO) mice characterised by: a significant increase in ear swelling (Figure 7E); significantly elevated vascular permeability quantified by Evans blue extravasation (Figure 7F); and enhanced tissue inflammation with increased numbers of degranulating mast cells revealed by Toluidine blue stained histopathology (Figure 7G). These findings establish mast cell steroidogenesis as a critical endogenous mechanism that maintains immune homeostasis by limiting the magnitude of inflammatory responses.

## DISCUSSION

Our study unveils a previously unrecognized role for de novo steroidogenesis in mast cell biology, revealing a primitive form of immune regulation that predates the evolution of adaptive immunity. Through an integrative multi-omics approach combined with functional assays, we demonstrate that mast cell-derived steroids are essential for their maturation, survival, functional regulation, and recovery after activation. These findings not only advance our understanding of mast cell physiology but also provide a foundation for developing targeted therapies for mast cell-associated pathologies.

### Mast cell steroidogenesis represents a primitive form of steroidogenesis

The discovery that mast cells express functional Cyp11a1 and produce pregnenolone represents a significant evolutionary insight. The evolution of glandular steroidogenesis shaped vertebrate evolution (Baker, 2019), including immune regulation. The most primitive step of steroidogenesis is the conversion of cholesterol to pregnenolone (Fan and Papadopoulos, 2013). The most primitive form of glandular steroidogenesis evolved in teleost (fishes). However, mast cells evolved an estimated 250 million years earlier in tunicates (St John et al., 2023). Our observation of mast cell steroidogenesis, particularly the conversion of cholesterol to pregnenolone, suggests that mast cells adapted this molecular mechanism earlier than the evolution of glandular steroidogenesis. This molecular adaptation likely evolved to regulate immune cell reactions and restore immune and tissue homeostasis. The immunoregulatory mechanism of mast cell steroidogenesis thus represents a primitive form of immune regulation long before the emergence of centralized endocrine organs and systemic steroid hormone-mediated immune regulation.

The presence of steroidogenic mast cells in adrenal glands and gonads, as revealed in this study, raises intriguing questions about their potential contribution to glandular steroidogenesis and the evolutionary relationship between immune and endocrine systems. To further elucidate the evolutionary significance of this process, it would be valuable to study mast cell steroidogenesis in lower chordates. Such investigations could provide deeper insights into the evolution of this immunoregulatory mechanism and its role in the broader context of vertebrate endocrine and immune system.

### Cell-intrinsic steroidogenesis as a negative regulator of mast cell function

Our findings reveal that mast cell-derived steroids act as critical negative regulators of mast cell activation. Cyp11a1-deficient mast cells exhibited heightened degranulation in response to both IgE-dependent and -independent stimuli, leading to exaggerated inflammatory responses *in vivo*. This suggests that endogenous steroidogenesis serves as a counter-regulatory mechanism to fine-tune mast cell activation and prevent excessive inflammation. Importantly, this mechanism operates both in IgE-dependent and independent responses, providing a novel pathway for regulating mast cell function.

### Insights into the regranulation phase: A critical period for immune homeostasis

Mast cells are long-lived cells with the capacity for several rounds of activation and degranulation. Degranulation, the release of preformed granule content, orchestrates the inflammatory responses to raise immunity. It is expected that the differentially expressed mRNAs and proteins during the regranulation phase are involved in the restoration of immune homeostasis and tissue repair. For our comparative study, we selected the eight-hour post-degranulation timepoint as the regranulation phase. This decision was based on previous observations that the transcriptional burst initiates within two to four hours after degranulation, while the translational burst begins later, and granule formation completes in 24 hours (Burwen, 1982; Dvorak et al., 1988; Iskarpatyoti et al., 2022; Jayapal et al., 2006; Kuehn et al., 2010; Xiang et al., 2001). Our transcriptomic and proteomic analyses provide new insights into the molecular events during the regranulation phase, revealing extensive remodelling of gene and protein expression profiles. Notably, we observed increased Cyp11a1 expression during regranulation, accompanied by elevated pregnenolone production. The dysregulation of regranulation-associated genes in Cyp11a1-deficient mast cells underscores the importance of steroidogenesis in coordinating recovery and tissue repair processes. Future studies should investigate how mast cell-derived steroids interact with other regulatory pathways during regranulation to coordinate recovery and tissue repair.

### Sexual dimorphism in mast cell steroidogenesis

Our proteomic analysis uncovered significant gender-specific differences in protein expression profiles, particularly during the regranulation phase and upon *Cyp11a1* deletion. These findings provide a molecular basis for the well-documented sexual dimorphism in immune responses and highlight the importance of considering sex as a biological variable in mast cell research. Future studies should investigate how hormonal regulation interacts with mast cell-intrinsic steroidogenesis to shape immune responses in males and females.

### Transcription-translation uncoupling: A previously unknown feature of mast cells

A surprising finding from our study was the uncoupling between transcriptional and translational changes in mast cells. Comparative analyses revealed minimal overlap between differentially expressed genes (DEGs) and proteins (DEPs), suggesting that post-transcriptional mechanisms play a dominant role in shaping the mast cell proteome. This uncoupling may reflect the need for rapid protein synthesis during activation while maintaining long-term plasticity for subsequent responses. Such uncoupling has been reported in specialized cell types such as neurons, oocyte and spermatids but has not been previously described in mast cells. This phenomenon may be particularly relevant for mast cells, given their dual roles as sentinels for immediate immune responses, venom detoxification and mediators of tissue remodelling over longer timescales. Future studies should investigate the molecular mechanisms underlying this uncoupling, including the roles of RNA-binding proteins, microRNAs, and translational regulators.

### Endogenous steroidogenesis in mast cells as a therapeutic gateway for inflammation

Our discovery of Cyp11a1-dependent steroidogenesis in mast cells as an endogenous anti-inflammatory mechanism offers a potential paradigm shift in treating allergic and inflammatory disorders. While synthetic glucocorticoids remain the clinical gold standard, their broad immunosuppressive effects underscore the need for targeted alternatives. Mast cell steroidogenesis represents a cell-autonomous regulatory axis that could be exploited for more precise therapeutic interventions.

Developing structurally optimised pregnenolone derivatives to amplify mast cell-specific steroid signalling could potentially mitigate the systemic side effects of exogenous glucocorticoids while maintaining or enhancing anti-inflammatory efficacy. This approach aligns with precision medicine principles, leveraging cell-type-specific pathways to minimize off-target effects. Moreover, the observed sex-specific differences in steroidogenic output highlight the potential for sex-stratified therapies, addressing the often-overlooked influence of patient sex on disease severity and treatment response in allergic and autoimmune conditions.

By integrating mast cell biology, steroid immunometabolism, and sex dimorphism, this work redefines therapeutic targeting in inflammation. It positions mast cell steroidogenesis as a druggable axis capable of recalibrating immune homeostasis with unprecedented specificity, potentially catalysing the development of smart anti-inflammatory agents that offer context-aware, pathology-restricted interventions.

In conclusion, our study reveals mast cell steroidogenesis as a fundamental regulatory mechanism in immune function, with far-reaching implications for our understanding of mast cell biology, evolution of immune regulation, and development of novel therapeutic strategies. The comprehensive datasets generated in this study provide a valuable resource for future investigations into mast cell heterogeneity, function, and their roles in health and disease.

### Limitations of the study

While our integrated multi-omics approach reveals fundamental aspects of mast cell steroidogenesis, several considerations merit attention. Though well-established, the reliance on BMMC and PMC models may not fully mirror the functional complexity of tissue-resident mast cells in their native microenvironments. Additionally, our murine model findings require validation in human systems to assess translational relevance. While we comprehensively characterized cell-intrinsic steroidogenic regulation, potential cell-extrinsic interactions between mast cell-derived steroids and other cell types such as immune, stromal, or any other cell types of inflamed tissue remain unexplored. Finally, the precise molecular mechanisms underlying pregnenolone’s immunomodulatory effects - including potential receptor interactions or non-genomic signalling pathways - represent an important area for future investigation. These considerations highlight opportunities to expand our findings across model systems and mechanistic scales while preserving the biological relevance of this primitive regulatory pathway.

### Resource availability

All data and code used in the analysis will be available in a public database before or immediately after the manuscript is accepted in principle. Additionally, we are creating a database with a user-friendly interface for browsing the omics data, which will be available before the revised submission of the manuscript.

## MATERIALS AND METHODS

### Mice

In this investigation, all mice were handled and cared for following the stringent guidelines set forth by the UK Animals in Science Regulation Unit’s Code of Practice for the Housing and Care of Animals Bred, Supplied, or Used for Scientific Purposes, and the Animals (Scientific Procedures) Act 1986 Amendment Regulations 2012. All experimental protocols were conducted under the authorization of a UK Home Office Project License (PPL PP2448972) and received the necessary approval from the local institute’s Animal Welfare and Ethical Review Body, ensuring compliance with ethical standards and animal welfare considerations. The sample size was determined according to our previous experience and a priori power analysis (G*Power). Housing condition of Gurdon animal facility: all mice used in this study were maintained in a specific pathogen-free unit on 12 hours of light and 12 hours of dark cycles. The mice were genotyped by Transnetyx. *Cyp11a1-*mCherry reporter and *Cyp11a1*^fl/fl^ mice were generated by Sanger Institute as previously described in Mahata et al., 2020 (Mahata et al., 2020). Mast cell & basophil specific knockout mice, *Cyp11a1*^fl/fl^;*Cpa3*^Cre^ were generated with crossing with *Cpa3-*Cre mice (Jackson laboratory). In this study we used mice aged between 8 to 12 weeks.

### Bone marrow-derived mast Cell (BMMC) differentiation and culture

Bone marrow cells were isolated from femurs and tibias of 6-12 week-old mice and cultured in RPMI 1640 medium supplemented with 10% FBS, 100 U/mL penicillin, 100 μg/mL streptomycin, 2 mM L-glutamine, 50 μM 2-mercaptoethanol, 30 ng/mL IL3, and 10 ng/mL SCF. Cells were incubated at 37°C, 5% CO2 for 4-6 weeks. Media was changed on 2^nd^ and 4^th^ day, and cells were transferred to a new flask followed by regular media change (every 5-7 days). Mast cell purity (>95%) was confirmed by flow cytometry analysis of FcεRI and c-Kit expression.

### Peritoneal mast cells (PMC) isolation and culture

We used 8 to 12-week-old female C57BL/6N mice. After culling, 7 ml of ice-cold sterile Tyrode’s buffer (Thermo Scientific, Cat. No. J67607.K2) and 3 ml of air were injected in the peritoneal cavity using a 10 ml syringe equipped with a 27 G needle. After injection, we shook the mice for 1 min to detach peritoneal cells into the Tyrode’s buffer. We aspirated the fluid from the abdomen gently and slowly (∼0.5 ml/s) to avoid clogging by the inner organs and transferred the collected cell suspension into a collection tube. Next, we centrifuged the tubes with the cell suspension at ∼300 x g for 7 mins at 4°C. Under a sterile hood, we aspirated the supernatant and washed cells with DMEM complete medium. We resuspended the pellets in 4 ml of pre-warmed DMEM complete medium with IL-3 (20 ng/ml) and Stem Cell Factor (SCF, 30 ng/ml), transferred the cell suspension to a 25 cm^2^ culture flask and incubated for 2-4 weeks (37 °C and 5% CO2). On 2nd and 4th day after plating cells, we changed the medium, and transferred the cell suspension on to a new culture flask and continued the culture for 2-4 weeks. We analysed their maturation (cKit and FceR1 expression) by flow cytometry.

### Mast cell degranulation assay

Mast cell degranulation was assessed by measuring β-hexosaminidase release as previously described (Tsvilovskyy et al., 2018), with modifications. BMMCs and PMCs were sensitized overnight with 1 μg/mL anti-DNP IgE, washed, and resuspended in Tyrode’s buffer at a concentration of 1 × 10^6^ cells/mL. Cells were then stimulated with DNP-HSA (10-100 ng/mL). Compound 48/80 (10 μg/mL) was used for IgE-independent degranulation for 30 minutes in Tyrode buffer. After stimulation, cell suspensions were centrifuged and placed on ice, and supernatants were collected. The cell pellets of unstimulated cells were lysed with 0.5% Triton X-100 to determine the maximal enzymatic activity of β-hexosaminidase. Twenty-five microliters of supernatant or lysate volume corresponding to 5 × 10^3^ cells was incubated with 50 μL of a 1.3 mg/mL p-nitrophenyl-N-acetyl-β-D-glucosaminide (aka pNAG) solution in 50 mmol/L citrate buffer pH 4.5 at 37°C for 90 minutes. The reaction was stopped with 150 μL of 0.2 mol/L glycine buffer, pH 10.7, and absorbance read at 405 nm. Calculate the percentage of β-hexosaminidase activities present in the supernatants. Percentage degranulation = 100 × (supernatant content)/(supernatant + lysate content) (Kuehn et al., 2010).

### Spectral Flow Cytometry

We followed eBioscience surface staining, intracellular cytotoplasmic protein staining (for cytokines) protocols. Briefly, single-cell suspensions were stained with Live/Dead Fixable Dead cell stain kit (Molecular Probes/ Thermo Fisher) and blocked by purified rat anti-mouse CD16/CD32 purchased from BD Bioscience and eBioscience. Surface staining was performed in flow cytometry staining buffer (eBioscience) or in PBS containing 3% FCS at 4 °C. After staining, cells were washed with flow cytometry staining buffer (eBioscience) or 3% PBS-FCS, and were analysed by Cytek Aurora (5L) flow cytometer. FlowJo v10.2 software facilitated the data analysis.

### Steroid quantification via liquid chromatography mass spectrometry (LC-MS/MS)

Samples (100 μL) of each cell supernatant sample, were enriched with isotopically labelled internal standards, including 13C2,d2-pregnenolone (1 ng) and extracted along with a mixed steroid calibration curve, including pregnenolone (0.005-1 ng) through supported liquid extraction plates on an Extrahera liquid handling robot (Biotage, Uppsala, Sweden) using dichloromethane/isopropanol (98:2 v/v), reduced to dryness under nitrogen and resuspension in water/methanol (80 μL; 70:30 v/v water/methanol) followed by LC-MS/MS analysis of the extract.

Frozen tissue (∼50 mg - exact weight recorded) was placed in 2 mL reinforced tubes (containing 1.4 mm ceramic beads, FisherScientific). Acetonitrile with 0.1 % formic acid (v/v; 1 mL) was added to the tube, and it was enriched with 20 uL isotopically labelled steroid standard mixture (as above). Each tube was added to a Bead Ruptor 24 Elite (Omni International) fitted with a CryoCool unit. The tubes were homogenised for 1 m/s for 30 seconds for 3 cycles. The supernatant for each sample was transferred to a Filter+ plate (Biotage, Sweden), positive pressure was applied, and the eluate was collected in a clean 96-well collection plate. The filtered homogenate was further processed through a phospholipid depletion (PLD+) plate (Biotage, Sweden) and the eluate was reduced to dryness, resuspended in water/methanol (70:30 v/v) and the plate sealed with a zone-free plate seal, ready for LC-MS/MS analysis.

Briefly, an I-Class UPLC (Waters, UK) was used for the liquid chromatography on a Kinetex C18 column (150 x 2.1 mm; 2.6 μm) with a flow rate of 0.3 mL/min and a mobile phase system of water with 0.05 mM ammonium fluoride and methanol with 0.05 mM ammonium fluoride, starting at 50% B, rising to 95% B and returning to 50% B. Separation of 18 steroids was carried out. The column and autosampler temperatures were maintained at 50 and 10℃, respectively. The injection volume was 20 µL and the total analytical run time per sample was 16 min. Steroids were detected on a QTrap 6500+ mass spectrometer (AB Sciex, Warrington, UK) equipped with an electrospray ionisation turbo V ion spray source. The positive ion spray voltage was set to 5500 V and the negative ion spray voltage was set to -4500 V, with the source temperature maintained at 600oC. Multiple reaction monitoring parameters were carried out for all steroids including pregnenolone (P5) m/z 317.1 281.1 and 159.0 with declustering potential (DP) of 66 collision exit potential (CXP) of 31 and 29 V and collision energy (CE) of 12 V, respectively and for 13C2,d2-pregnenolone of 321.2 * 285.2 with DP of 14 CXP of 17 and CE of 18 with retention time of 10.4 mins.

The ratio of P5/13C2,d2-P5 peak areas were calculated and linear regression analysis used to calculate the amount of P5 in each sample. The same was done for other steroids in the sample (aldosterone, progesterone, 17β-estradiol, 5α-dihydrotestosterone and testosterone) by evaluation of the data on MultiQuant 3.0.3 (AB Sciex, UK).

### RNA-seq analysis

Quality assessment of the raw sequencing reads was conducted using FastQC, and the HISAT2 aligner was utilized to map the reads to the human genome (hg38), resulting in SAM files for each sample. Samtools was subsequently used to convert and sort these files into BAM format. Gene-level abundance and raw counts were derived using htseq-count, providing the necessary data for principal component analysis (PCA) and differential gene expression analysis in DESeq2 (version 1.30.1) (Love et al., 2014). Gene set enrichment analysis (GSEA) (Subramanian et al., 2005) were performed with clusterProfiler (version 3.18.1) (Yu et al., 2012).

To assign immune cell phenotypes within the TME for deconvolution analysis, the CIBERSORTx web portal (https://cibersortx.stanford.edu/runcibersortx.php) was utilized. The gene expression matrix derived from TNBC patients was inputted into the CIBERSORT tool. For this analysis, the LM22 signature matrix, which includes gene expression profiles for established immune markers, was employed as the default cell-type signature matrix. Correlations and p-values were calculated to explore the relationships between each steroid hormone receptor genes and the different immune population scores derived from CIBERSORT. This involved a systematic iteration over the selected genes, during which the Pearson correlation coefficient was computed for each gene in relation to the immune cell score. To assess the statistical significance of these correlations, p-values were determined. Following this, adjustments were made to the p-values to ensure accurate interpretation of significance levels and bolster the reliability of the results.

### Passive cutaneous anaphylaxis (PCA) reaction in mice ear

6-8 week old *Cpa3*^Cre^ (control) or Cyp11a1 knock-out mice (*Cyp11a1*^fl/fl^;*Cpa3*^Cre^) mice were sensitized by intradermal injection of 0.4 μg anti-DNP IgE in 10 μL PBS or PBS only into the ear. After 24 hours, mice were challenged intravenously via the tail vein with 200 μL of a solution containing 10 μg DNP-HSA and 2% Evans blue dye in PBS. Ear thickness was measured using a digital calliper immediately before and 1 hour after the challenge. One hour post-challenge, mice were euthanized, and ears were excised. Each ear was homogenized in 1 mL PBS, followed by the addition of acetone (3:7 v/v). After overnight incubation at room temperature, samples were centrifuged at 3000 rpm for 15 minutes. The absorbance of the supernatant was measured at 620 nm to quantify Evans blue as a surrogate of plasma extravasation.

### Toluidine blue staining

Tissue samples from mouse ears, tongue, and caecum were fixed in 10% neutral buffered formalin for 24 hours at room temperature. Fixed tissues were dehydrated through a graded ethanol series, cleared in xylene, and embedded in paraffin. Sections of 4-8 μm thickness were cut using a microtome and mounted on glass slides. For Toluidine Blue staining, sections were deparaffinized in xylene and rehydrated through a graded ethanol series. Slides were then stained with 0.1% Toluidine Blue O solution (pH 2.0-2.5) for 5-10 minutes at room temperature. After rinsing in distilled water, sections were rapidly dehydrated through 95% and 100% ethanol, cleared in xylene, and mounted with a coverslip using a permanent mounting medium. This method results in metachromatic staining of mast cell granules, appearing purple-red against a blue background. Mast cells were quantified by counting Toluidine Blue-positive cells in multiple high-power fields per section using light microscopy. We used NanoZoomer to capture images and figures made using NDP.view2 software.

### Sample dissolution, TMT labelling and reverse-phase fractionation for proteomics

Tissue samples were resuspended in lysis buffer containing 100mM Triethylammonium bicarbonate (TEAB, Sigma), 10% Isopropanol, 50mM NaCl, 1% SDC with Nuclease and Protease/Phosphatase Inhibitors, followed by 15 min incubation at RT. Protein concentration was estimated using Bradford assay according to manufacturer’s instructions (BIO-RAD-Quick start). Total proteins were reduced and alkylated with 2ul of 50mM tris-2-caraboxymethyl phosphine (TCEP, Sigma) and 1ul of 200mM Iodoacetamide (IAA, Sigma) respectively for 1 hour. Then protein samples were digested overnight at 37°C using trypsin solution at ratio protein/trypsin ∼ 1:30. The next day, protein digest was labelled with the TMTpro reagents (Thermo Scientific) for 1 hour and quenched with 5% hydroxylamine (Thermo Scientific) for 15 min at room temperature (RT). All the samples were mixed and acidified using Formic Acid for SDC removal. The dry TMT mix was fractionated on a Dionex Ultimate 3000 system at high pH using the X-Bridge C18 column (3.5 μm, 2.1x150mm, Waters) with 90min linear gradient from 5% to 95% acetonitrile contained 20mM ammonium hydroxide at a flow rate of 0.2 ml/min. Peptides fractions were collected between 20-55 minutes and were dried with speed vac concentrator. The fractions were reconstituted in 0.1% formic acid for liquid chromatography tandem mass spectrometry (LC–MS/MS) analysis (Mohammed et al., 2016).

#### LC-MS/MS

Peptide fractions were analysed on a Dionex Ultimate 3000 system coupled with the nano-ESI source Fusion Lumos Orbitrap Mass Spectrometer (Thermo Scientific). Peptides were trapped on a 100μm ID X 2 cm microcapillary C18 column (5µm, 100A) followed by 2h elution using 75μm ID X 25 cm C18 RP column (3µm, 100A) at 300nl/min flow rate. In each data collection cycle, one full MS scan (380– 1,500 m/z) was acquired in the Orbitrap (120K resolution, automatic gain control (AGC) setting of 3×105 and Maximum Injection Time (MIT) of 100 ms). The subsequent MS2 was conducted with a top speed approach using a 3-s duration. The most abundant ions were selected for fragmentation by collision induced dissociation (CID). CID was performed with a collision energy of 35%, an AGC setting of 1×104, an isolation window of 0.7 Da, a MIT of 35ms. Previously analysed precursor ions were dynamically excluded for 45s. During the MS3 analyses for TMT quantification, precursor ion selection was based on the previous MS2 scan and isolated using a 2.0Da m/z window. MS2–MS3 was conducted using sequential precursor selection (SPS) methodology with the top10 settings. HCD was used for the MS3, it was performed using 55% collision energy and reporter ions were detected using the Orbitrap (50K resolution, an AGC setting of 5×104 and MIT of 86 ms).

Phospho fractions were subjected to MS2 analysis only.

#### Data processing

The Proteome Discoverer 3.0. (Thermo Scientific) was used for the processing of CID tandem mass spectra. The SequestHT search engine was used and all the spectra searched against the Uniprot Homo sapiens FASTA database (taxon ID 9606 - Version June 2023). All searches were performed using a static modification of TMTpro (+304.207 Da) at any N-terminus and lysines and Carbamidomethyl at Cysteines (+57.021 Da). Methionine oxidation (+15.9949Da), Phospho (+79.966 Da on Serine, Threonine and Tyrosine) and Deamidation on Asparagine and Glutamine (+0.984) were included as dynamic modifications. Mass spectra were searched using precursor ion tolerance 20 ppm and fragment ion tolerance 0.5 Da. For peptide confidence, 1% FDR was applied, and peptides uniquely matched to a protein were used for quantification.

Analysis was carried out using the qPLEXanalyzer package (version1.16.1) in R (Papachristou et al., 2018).

### Quantitative Pregnenolone ELISA

Pregnenolone concentrations of the cell culture supernatants were quantified using a pregnenolone ELISA kit (Abnova) following the manufacturers’ instructions. Absorbance was measured at 450 nm, and data were analyzed in GraphPad Prism 6.

### Statistics

For the calculation of DEGs in bulk RNA-seq datasets, p-values were derived using the default methodologies embedded in the respective R packages. The ggpubr R package (version 0.4.0), accessible at (https://CRAN.R-project.org/package=ggpubr), was employed to compare differences in specific gene set scores and metabolic flux levels across immune cells from varying groups.

In both in vitro and in vivo experimental settings, Prism9 (GraphPad Software) was utilized to ascertain the significance levels of the data, applying a two-tailed Student’s t-test. A p-value falling below the 0.05 threshold was deemed to indicate statistical significance.

## Acknowledgements

We would like to thank Dr Timotheus YF Halim, CRUK Cambridge Institute, for his help and support with cutaneous anaphylaxis experiments; Prof Gunnar Pejler and Mirjan Grujic for their helpful suggestions; Joana Cerveira, Cytometry facility, Dept. of Pathology and Richard Grenfell, CRUK CI Cytometry facility, for help with flow cytometry; Lorna Roberts, Histology facility; Clive D’Santos, Valar Nila Roamio Franklin, Abigail Edwards, CRUK CI Proteomics facility; UBS animal facility, Gurdon Institute, for their technical help and animal husbandry. We used BioRender.com to generate the graphical illustrations presented in this manuscript. The Figure was partly generated using Servier Medical Art, licensed under a Creative Commons Attribution 3.0 unported license AI.

## Author contribution

JP: Conceptualised, designed and performed experiments. Analysed, assembled, and visualised data. Wrote the manuscript. QZ: Performed all bioinformatics analyses, assembled and visualised data. NH: Involved with optimisation of samples, steroid detection, and mass-spectrometry data analysis. YYK: Performed intravenous injections. HH, SC and SKS: Were involved in tissue processing. RR and KO: Supported *in vivo* aspects of the study. BM: Led the project. Supervised the study. Reviewed the manuscript. Involved with conceptualization, experimental designing, fund acquisition and resource management. All authors read and approved the draft manuscript before submission.

## Data and materials availability

All data and code used in the analysis will be available in a public database before or immediately after acceptance (in principle) of the paper.

---

**Supplementary Figure S1.**
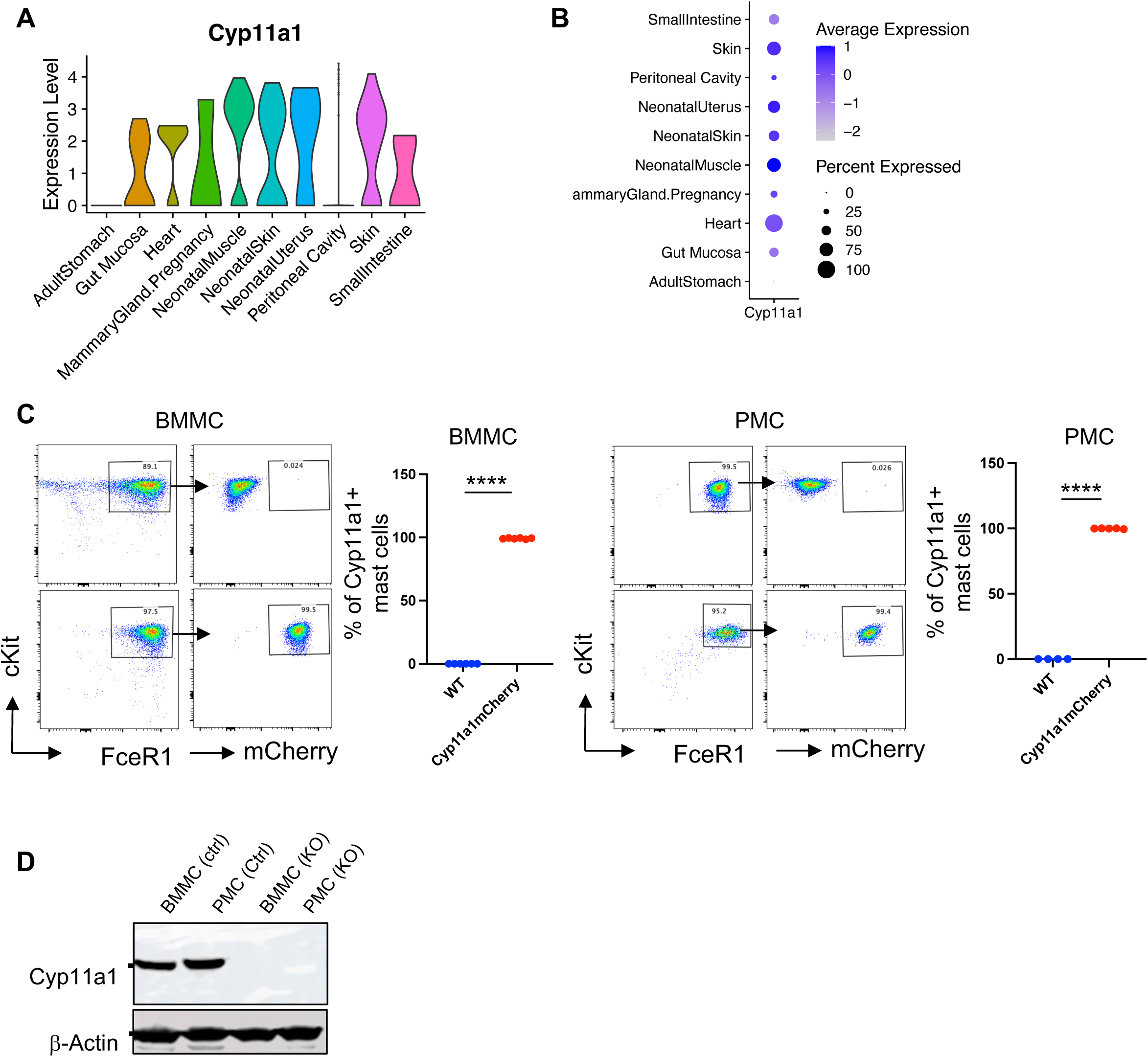
Analysis of Cyp11a1 expression and steroidogenic capacity in mast cells. **A.** Violin plots showing *Cyp11a1* expression in mast cells across diverse mouse tissues. Single-cell RNA sequencing data from adult stomach, gut mucosa, heart, pregnant mammary gland, neonatal muscle, skin, uterus, peritoneal cavity, and small intestine were analysed. Plot width indicates cell density at each expression level. RNAseq data were obtained from public repositories (see Methods). **B.** Dot plot representation of *Cyp11a1* expression in tissue-resident mast cells. Dot size represents the proportion of *Cyp11a1*-expressing cells; colour intensity indicates mean expression levels. Analysis corresponds to tissues shown in (A). **C.** Flow cytometric analysis of Cyp11a1 expression in bone marrow-derived (BMMC) and peritoneal (PMC) mast cells. Cells were differentiated *in vitro* from Cyp11a1-mCherry reporter or wild-type (WT) mice. FACS plots (representative) and histograms show mCherry expression as a surrogate for Cyp11a1. Representative of four independent experiments. p-value was calculated using unpaired two-tailed t-test. **D.** Western blot analysis confirming *Cyp11a1* deletion efficiency in BMMCs and PMCs derived from *Cyp11a1*^fl/fl^ ;*Vav1*^cre^ (KO) and *Vav1*^cre^ control mice. β-Actin serves as loading control. Representative of three independent experiments. Representative of three independent experiments.

**Supplementary Figure S2.**
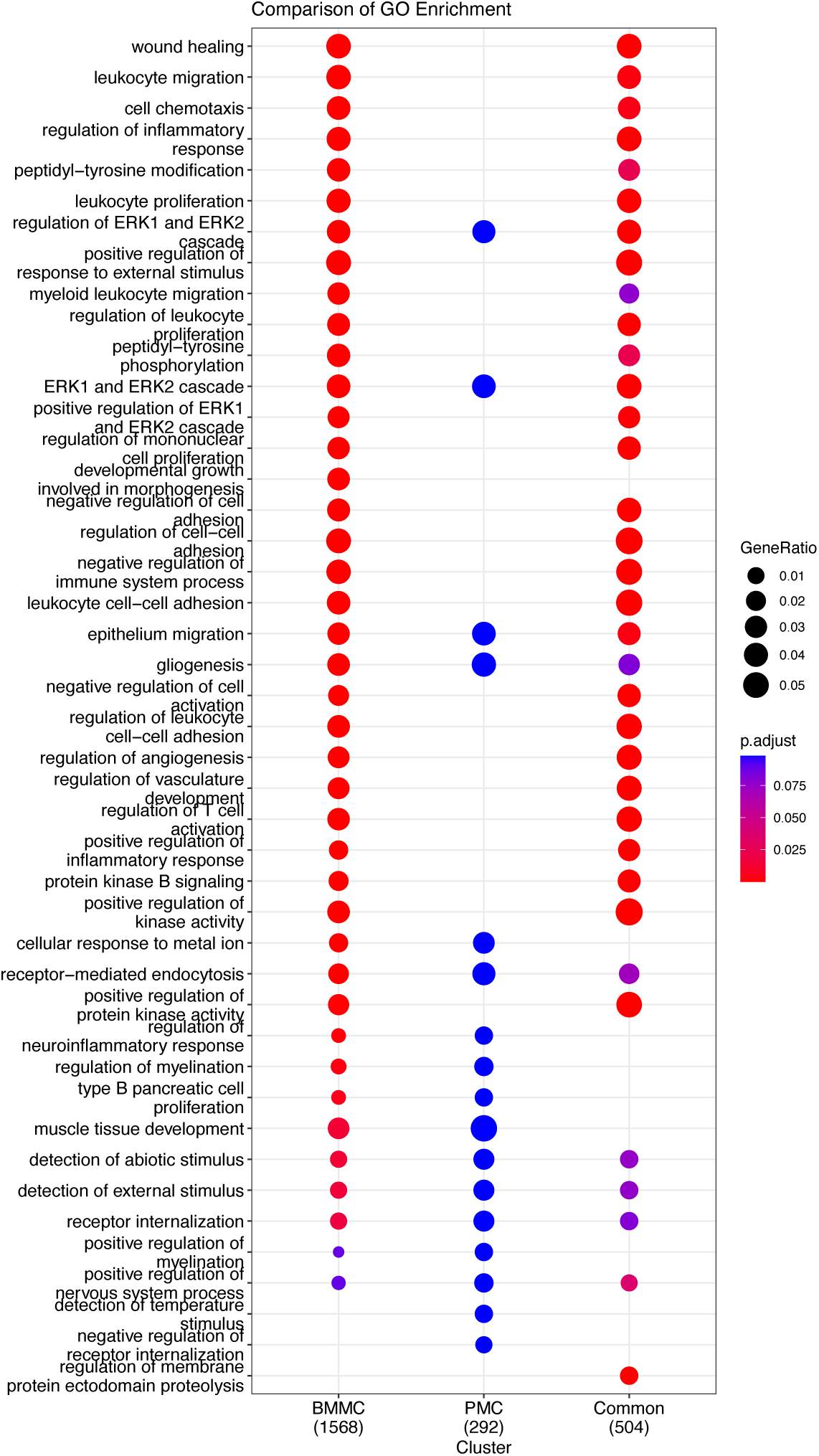
Functional commonness in differentially expressed genes during regranulation in BMMC and PMC. **A.** Visualisation of commonness in GO enriched pathways during regranulation compared to their mature state (as revealed in Figure 2H and I). The dot plot shows the GO enrichment among the unique DEGs from BMMC and PMC mature mast cells, and the common DEGs from both groups. Each dot represents a GO term, with the size of the dot indicating the gene ratio and the colour indicating the adjusted p-value for enrichment.

**Supplementary Figure S3.**
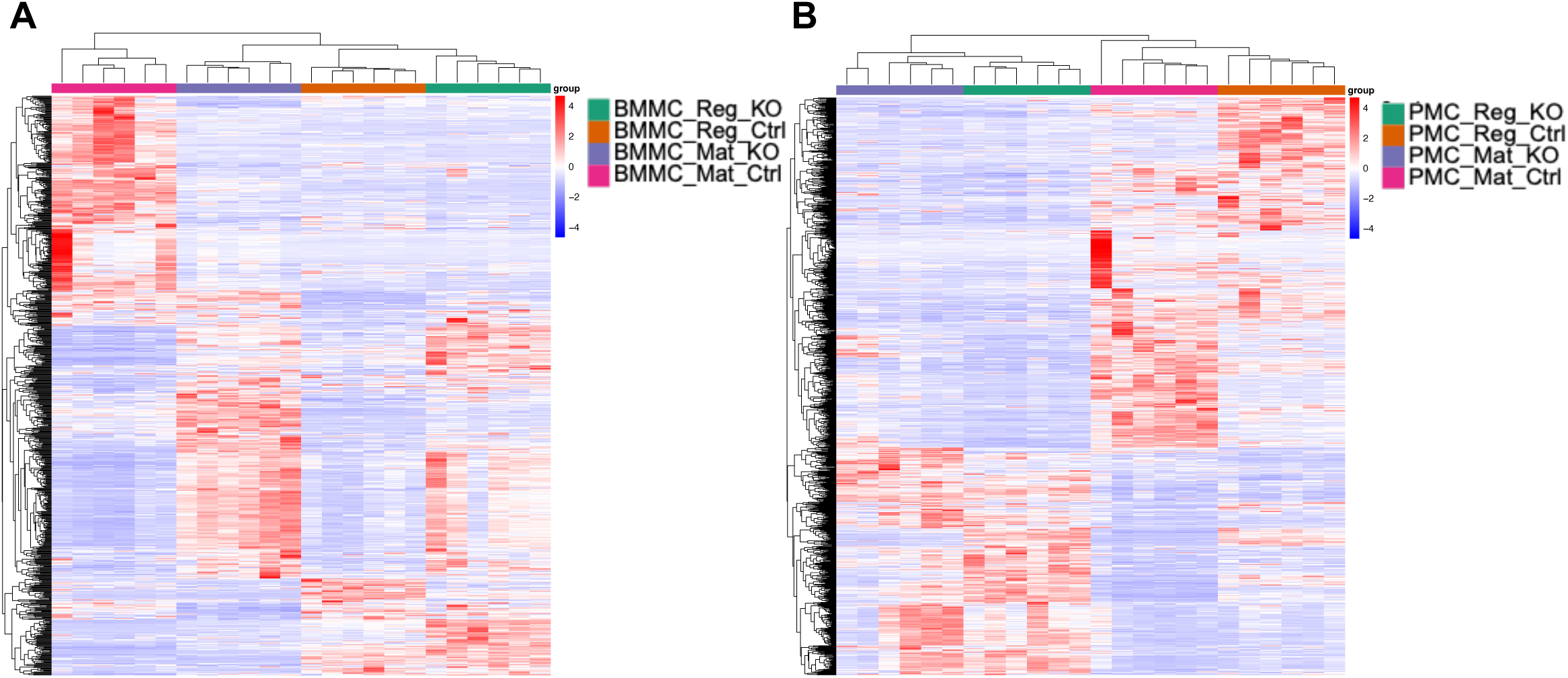
Effect of *Cyp11a1* deletion in mature and regranulating mast cells. **A-B.** The heatmaps show the expression of significant differentially expressed genes comparing Cyp11a1-deficient (KO) with Cyp11a1-sufficient control mast cells in all four BMMC (A) and PMC (B) groups.

**Supplementary Figure S4.**
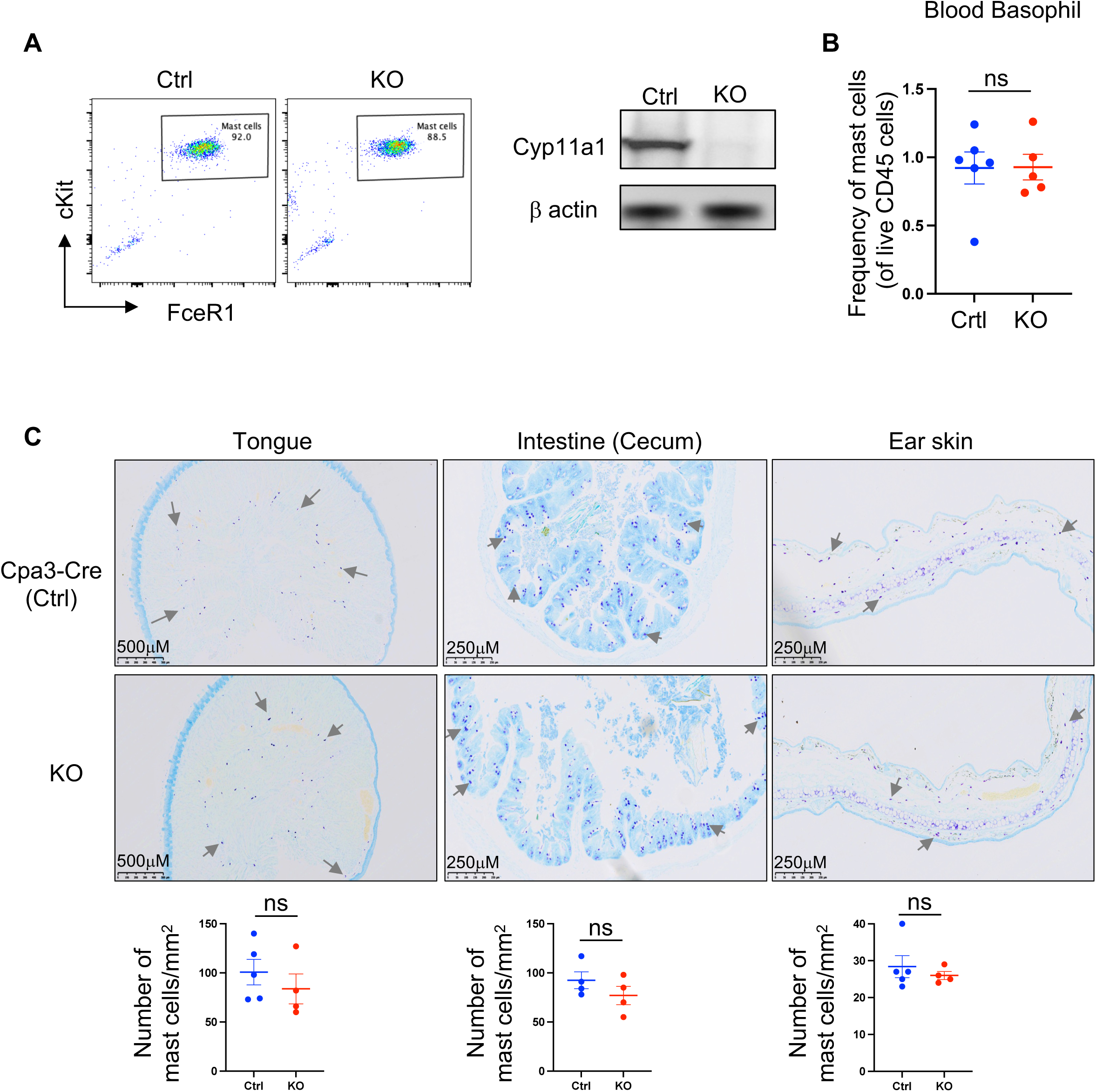
Effect of Cpa3-Cre mediated *Cyp11a1* knockout in *Cyp11a1*^fl/fl^;*Cpa3*^Cre^ (KO) mice on mast cell and basophil development and tissue distribution. **A.** Peritoneal cavity-derived mast cells (PMC) were differentiated in vitro and analysed for their maturation by cKit and FceR1 expression (left-panel, Flow Cytometry, representative). PMCs were used in western blot analysis to detect Cyp11a1 deletion efficiency (right-panel). Cpa3-Cre mice were used as a control to show Cyp11a1. β-actin was used as a loading control in the western blot. **B.** The frequency of blood basophils in control (*Cpa3*^Cre^) and *Cyp11a1*^fl/fl^;*Cpa3*^Cre^ (KO) mice was measured by flow cytometry. Gating: All cells > Singlets > live CD45^+^ cells > Lineage^-^ > cKit^-^FceR1^+^ **C.** Toluidine blue staining of tissue sections derived from control (*Cpa3*^Cre^) and *Cyp11a1*^fl/fl^;*Cpa3*^Cre^ (KO) mice. Respective quantifications of mast cells have been shown in the bottom row. All experiments are representative of three independent experiments. p-value was calculated using unpaired two-tailed t-test

